# Evolutionary dynamics of abundant stop codon readthrough in *Anopheles* and *Drosophila*

**DOI:** 10.1101/051557

**Authors:** Irwin Jungreis, Clara S Chan, Robert M Waterhouse, Gabriel Fields, Michael F Lin, Manolis Kellis

**Author notes:** These authors contributed equally to this work (Irwin Jungreis, Clara S Chan).

## Abstract

Translational stop codon readthrough was virtually unknown in eukaryotic genomes until recent developments in comparative genomics and new experimental techniques revealed evidence of readthrough in hundreds of fly genes and several human, worm, and yeast genes. Here, we use the genomes of 21 species of *Anopheles* mosquitoes and improved comparative techniques to identify evolutionary signatures of conserved, functional readthrough of 353 stop codons in the malaria vector, *Anopheles gambiae*, and 51 additional *Drosophila melanogaster* stop codons, with several cases of double and triple readthrough including readthrough of two adjacent stop codons, supporting our earlier prediction of abundant readthrough in pancrustacea genomes. Comparisons between *Anopheles* and *Drosophila* allow us to transcend the static picture provided by single-clade analysis to explore the evolutionary dynamics of abundant readthrough. We find that most differences between the readthrough repertoires of the two species are due to readthrough gain or loss in existing genes, rather than to birth of new genes or to gene death; that RNA structures are sometimes gained or lost while readthrough persists; and that readthrough is more likely to be lost at TAA and TAG stop codons. We also determine which characteristic properties of readthrough predate readthrough and which are clade-specific. We estimate that there are more than 600 functional readthrough stop codons in *A. gambiae* and 900 in *D. melanogaster*. We find evidence that readthrough is used to regulate peroxisomal targeting in two genes. Finally, we use the sequenced centipede genome to refine the phylogenetic extent of abundant readthrough.

## Introduction

Although a ribosome will normally terminate translation when it encounters one of the three stop codons, UAG, UGA, and UAA, it will sometimes instead insert an amino acid and continue translation in the same frame, adding a peptide extension to that instance of the protein, a phenomenon known as stop codon readthrough (Doronina and Brown 2006; Namy and Rousset 2010). The tRNA that inserts the amino acid at the stop codon can be a selenocysteine tRNA if there is a downstream selenocysteine insertion sequence (SECIS element), a cognate of the stop codon in organisms that contain such “stop suppressor” tRNAs, or a near cognate tRNA that inserts its cognate amino acid with some frequency at certain “leaky” stop codons (Bonetti et al. 1995; Poole et al. 1998; Blanchet et al. 2014). The rate of leakage can depend on the choice of stop codon, the immediate stop codon context, particularly the 3’ nucleotide (Cridge et al. 2006; Brown et al. 1990b, 1990a; Loughran et al. 2014; Dabrowski et al. 2015), the presence of RNA structures in the mRNA (Steneberg and Samakovlis 2001; Brown et al. 1996; Firth et al. 2011; Wills et al. 1991; Houck-Loomis et al. 2011; Hirosawa-Takamori et al. 2009), trans factors within the cell (von der Haar and Tuite 2007; Beznosková et al. 2015), oxygen and glucose deprivation (Andreev et al. 2015), hydroxylation of the ribosomal decoding center (Loenarz et al. 2014), and other conditions that are not well understood. Readthrough has been proposed as an evolutionary catalyst in yeast, where both readthrough and frameshifting are epigenetically controlled via a prion protein state, thus enabling the adaptation of new domains translated at low rates during normal growth but at higher rates in periods of stress when they might provide a selective advantage (True and Lindquist 2000; Baudin-Baillieu et al. 2014).

Readthrough is common in viruses, where it increases functional versatility in a compact genome and provides a way to control the ratio of two protein isoforms (Namy and Rousset 2010). On the other hand, until recently only a handful of eukaryotic wild-type genes were known to exhibit readthrough (Klagges et al. 1996; Robinson and Cooley 1997; Steneberg and Samakovlis 2001; Namy et al. 2002, 2003; True and Lindquist 2000).

The first indication that readthrough was more prevalent in eukaryotic genomes came when the evolutionary lens of comparative genomics was turned upon 12 *Drosophila* genomes (Drosophila 12 Genomes Consortium et al. 2007; Stark et al. 2007). The pattern of substitutions provides an evolutionary signature that distinguishes protein-coding regions from non-coding ones, and continuation of this pattern beyond a stop codon until the next in-frame stop codon is an indication of conserved stop codon readthrough (Figure 1). A search for this evolutionary signature of readthrough identified 149 *Drosophila melanogaster* candidate readthrough transcripts, suggesting not only that translation does not always stop at the stop codon but also that the specific polypeptide sequence of the extended protein confers selective advantages at the protein level (Lin et al. 2007). Continuing this work, we expanded the list of readthrough candidates to 283 in *D. melanogaster*, 4 in human, and 5 in *C. elegans*, using improved comparative methods; ruled out alternative explanations for the evolutionary signatures; and experimentally validated several of the candidates (Jungreis et al. 2011; Lindblad-Toh et al. 2011). These readthrough candidates differed as a group from most other transcripts regarding their 4-base stop codon context, stop codon conservation, presence of RNA structures, and many other properties. Intrigued by the nearly two orders of magnitude greater prevalence of readthrough transcripts in *Drosophila* versus human and *C. elegans*, a phenomenon we termed “abundant readthrough”, we developed a statistical test using k-mer frequencies downstream of the stop codon to estimate the number of readthrough transcripts in a species using a single genome. Applying that test to 25 eukaryotic species led us to conjecture that abundant readthrough was present in insects and crustacea, but not in species outside the Pancrustacea clade.

**Figure 1.**
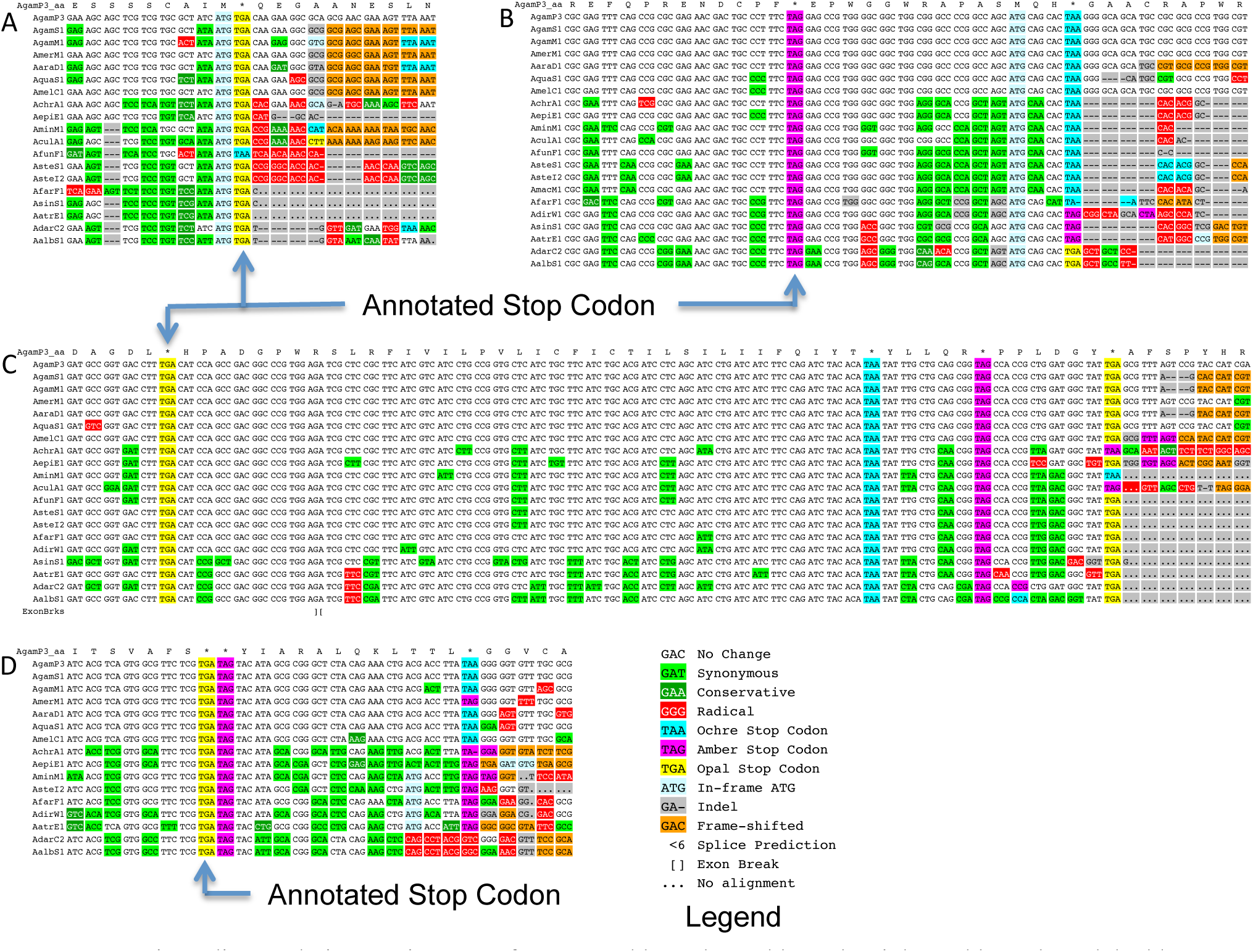
Protein-coding evolutionary signatures for non-readthrough, readthrough, triple readthrough, and double-stop readthrough stop codons. Alignments surrounding the annotated stop codons of four genes for 21 *Anopheles* species, displayed by CodAlignView (“CodAlignView: a tool for visualizing protein-coding constraint”, I Jungreis, M Lin, C. Chan, M Kellis, in preparation). The color coding of substitutions and insertions/deletions (indels) relative to *A. gambiae* is a simplification for visualization purposes, as the actual PhyloCSF score sums over all possible ancestral sequences and weighs every codon substitution by its probability. Insertions in other species relative to *A. gambiae* are not shown. (A) Alignment of a typical gene (AGAP011673-RA), shows abundant synonymous and conservative substitutions (green) upstream (to the left) of the stop codon, and many radical substitutions (red), frameshifting indels (orange), and poorly-conserved in-frame stop codons downstream of the annotated stop codon. The stop codon locus shows a substitution between different stop codons. (B) Alignment of AGAP000058-RA, one of 353 *A. gambiae* readthrough candidates. The region between the annotated stop codon and the next in-frame stop codon shows mostly synonymous substitutions and lacks frameshifting indels, while the region downstream from the second stop shows radical substitutions and indels typical of non-coding regions, providing evidence of continued protein-coding selection in the region between the two stop codons, and suggesting likely translational readthrough of the first stop codon. As is typical for readthrough candidates, the first stop codon is perfectly conserved, while the second stop codon shows substitutions between different stop codons. (C) Alignment of triple-readthrough candidate AGAP006474-RA (one of 35 double-readthrough candidates in *A. gambiae* including 5 triple-readthrough candidates). The second, third, and fourth ORFs all show protein-coding signatures, indicating that the first three stop codons are likely readthrough. None of these three stop codon positions show substitutions. (D) Alignment of double-stop readthrough candidate AGAP009063-RA (one of 13 cases). The ORF after two adjacent stop codons shows a protein-coding signature, indicating that the ribosome likely reads through both stop codons. To our knowledge, no cases of readthrough of two adjacent stop codons have previously been observed or predicted.

Since then, interest in readthrough in eukaryotes has blossomed. Readthrough has been demonstrated in human vascular endothelial growth factor A (VEGF-A), producing an isoform that reverses the angiogenic properties of VEGF-A, is regulated by a ribosomal binding protein, and is suppressed in colon cancer cells, having direct relevance to cancer treatment (Eswarappa et al. 2014; Eswarappa and Fox 2015). Readthrough of human Myelin protein zero produces an extended protein, L-MPZ, that is localized in compact myelin and could be involved in myelination (Yamaguchi et al. 2012). Readthrough has been shown to add peptide extensions to the genes encoding the human LDHB and MDH1 enzymes and several yeast genes that target the protein to the peroxisome (Schueren et al. 2014; Stiebler et al. 2014; Freitag et al. 2012). Mutational studies have shown that readthrough in 4 human genes predicted by comparative methods is triggered by a UGA-CUAG motif at the stop codon, a motif also found in a readthrough stop codon of the chikungunya virus (Loughran et al. 2014). One of our *Drosophila* readthrough candidates was found to exhibit readthrough in a heterologous yeast system (Chan et al. 2013), whereas several candidates having predicted stem loops did not exhibit high levels of readthrough, suggesting that readthrough in these stem-loop containing candidates might be modulated by trans factors in their native species. Readthrough has been proposed as the mechanism by which EFLGa peptides are created in the annelid *Platynereis dumerilii* (Conzelmann et al. 2013).

Ribosome profiling provided another opportunity to study readthrough at the whole genome level in a cell type-and condition-specific way by sequencing ribosome-protected fragments of mRNAs (Ingolia et al. 2009; Brar and Weissman 2015; Legendre et al. 2015). Ribosome profiling experiments detected readthrough in 350 *D. melanogaster* transcripts (S2 and embryonic cells), 42 human transcripts (foreskin fibroblasts), and several *S. cerevisiae* transcripts ([psi-] cells), some with readthrough rates of more than 50% and with different rates in the two *Drosophila* cell types providing evidence of regulation (Dunn et al. 2013; Artieri and Fraser 2014). In addition to validating 43 of our *Drosophila* readthrough candidates in these cell types, ribosome profiling detected several hundred genes in which readthrough occurs but has left no detectable evolutionary signature across species, supporting a hypothesis that readthrough arises initially as random failure of translation termination, and then, if the protein extension provides a benefit, is molded by selection into a conserved readthrough event. Readthrough has also been proposed to explain ribosome footprints within the 3’ UTRs of several *Plasmodium falciparum* transcripts (Caro et al. 2014; Bunnik et al. 2013).

These developments highlight the importance of annotating and studying readthrough genes in diverse species, both because of the wide ranging biological functions of readthrough in the particular species in which it occurs and in order to better understand the phenomenon of readthrough itself. Many of the questions from our *Drosophila* readthrough study remain unanswered. In most cases, the function of the readthrough polypeptide extension, the mechanism of readthrough, and the regulation of readthrough remains a mystery, as do the full extent and causes of abundant readthrough. Finally, it is unknown if the properties of readthrough transcripts found in *Drosophila* are specific to that clade or more general features of readthrough.

Just as zooming out from the genome of a single *Drosophila* species to compare many related species within the genus provided a powerful perspective for understanding that genome, so too zooming further out to compare two clades at greater evolutionary divergence can yield further insights. While analysis of a single clade provided a static picture of readthrough, comparison of two clades can provide insight into the evolutionary dynamics of readthrough, which can help to resolve the unanswered questions about abundant readthrough. To this end, we took advantage of the sequencing of multiple genomes of *Anopheles* mosquitoes to apply our comparative approaches to catalog an initial set of readthrough candidates in the malaria vector, *A. gambiae* (Neafsey et al. 2015). Here we report the improved comparative techniques for distinguishing readthrough genes used to identify those candidates as well as several more in both *A. gambiae* and in *D. melanogaster*; use orthology between *Drosophila* and *Anopheles* to better understand the characteristic properties and evolution dynamics of readthrough in these two clades; and obtain more precise bounds on the extent of abundant readthrough, both within and across species.

## Results

### *Anopheles* and *Drosophila* readthrough candidates

We began by generating a list of annotated *A. gambiae* PEST-strain transcripts that show evolutionary evidence of translation 3’ of the stop codon and for which translational stop codon readthrough unrelated to selenocysteine insertion is a more likely explanation than any of the alternatives. Using 21-way whole genome alignments of *Anopheles* species (Neafsey et al. 2015), for each annotated protein-coding transcript we evaluated the coding potential of the region between the annotated stop codon (“first stop codon”) and the next in-frame stop codon (“second stop codon”), which we refer to as the “second open reading frame (ORF)”, or, if the stop codon is read through, as the “readthrough region”. We will refer to the annotated coding region as the “first ORF”. Our procedure built on the one we had used previously in *Drosophila (Jungreis et al. 2011)* with additional steps to identify a more comprehensive list of candidates, and is summarized in Figure 2A.

**Figure 2.**
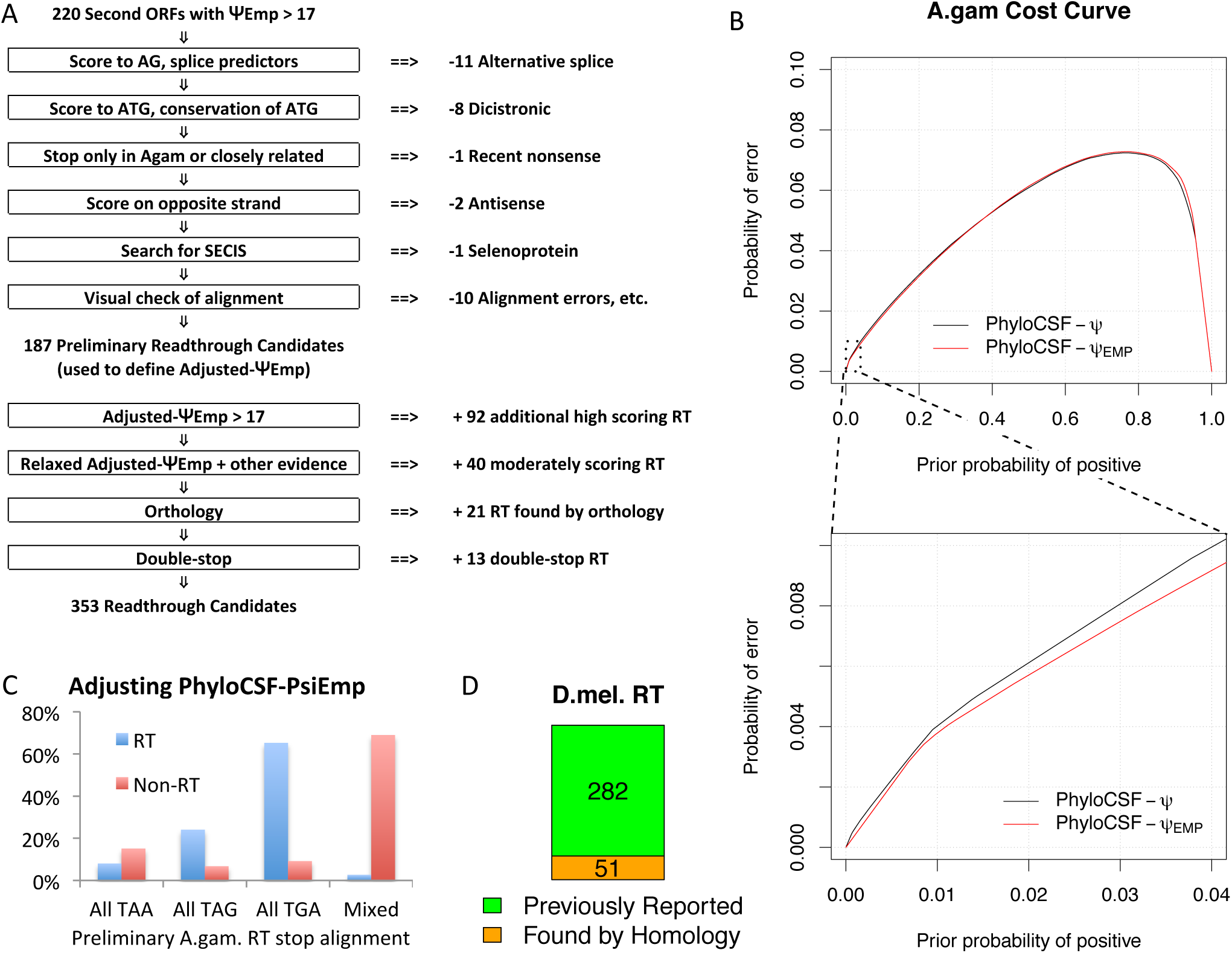
New comparative techniques and manual curation distinguishes 353 readthrough candidates in *A. gambiae* and 51 additional candidates in *D. melanogaster*. (A) Steps used to generate list of readthrough candidates in *A. gambiae*. Starting with 220 second ORFs having high PhyloCSF-Ψ_Emp_ score, we eliminated cases with a more plausible explanation of the protein-coding signature to yield 187 preliminary readthrough candidates. We used these to train Adjusted-Ψ_Emp_, and used that and other evidence to find 132 additional RT candidates. We then found 21 additional candidates using orthology to *D. melanogaster* candidates, and 13 by evaluating the ORFs downstream of two adjacent stop codons. (B) PhyloCSF-Ψ_Emp_ is a comparative method for distinguishing protein-coding regions that improves on our earlier method, PhyloCSF-Ψ, when extremely high specificity is required. Cross-validated cost curve (Drummon and Holte, 2000) shows, for each prior probability that the input region is coding, the probability that the discriminator makes an error, either false positive or false negative, at the optimal score threshold for that prior. The performance of PhyloCSF-Ψ (black curve) and of PhyloCSF-Ψ_Emp_ (red curve) are similar for most values of the prior (upper panel), but when the prior probability of coding is extremely low, PhyloCSF-Ψ_Emp_ makes noticeably fewer errors (lower panel). For example, PhyloCSF-Ψ_Emp_ makes approximately 7% fewer errors when the prior probability is 2%, corresponding to the fraction of D. melanogaster stop codons catalogued as readthrough in our earlier paper. (C) Data from preliminary readthrough candidates used to train Adjusted-Ψ_Emp_. Figure shows the fraction of preliminary readthrough candidate first stop codons (blue) and all other annotated stop codons (red) having an aligned stop codon in at least 10 species for which all aligned stop codons are TAA, all are TAG, all are TGA, or there is a mix of different aligned stop codons. For almost all preliminary readthrough candidates, the first stop codon is perfectly conserved, usually TGA, whereas the majority of other annotated stop codons are aligned to a mix of different stop codons. We used this information to define Adjusted-Ψ_Emp_ of a second ORF by determining to which of these four categories its first stop codon belongs, and combining that evidence with its PhyloCSF-Ψ_Emp_ score. (D) Readthrough candidates in *D. melanogaster*. For our comparative analyses, we used 333 *D. melanogaster* readthrough candidates consisting of 282 that had been reported in our earlier paper and 51 newly reported readthrough candidates found by homology to our *A. gambiae* candidates or the other *D. melanogaster* candidates.

We scored the protein-coding potential of each second ORF using PhyloCSF-Ψ_Emp_ a new variant of PhyloCSF-Ψ that is particularly good at excluding non-coding false positives in order to identify the small number of readthrough needles in the large haystack of second ORFs (Figure 2B and Methods). In brief, PhyloCSF is a comparative method that uses substitutions and codon frequencies to detect functional, conserved, protein-coding regions of genomes, while PhyloCSF-Ψ is a variant of PhyloCSF that accounts for the correlation between nearby codons by approximating the distribution of PhyloCSF scores on coding and non-coding regions with a family of normal distributions (Lin et al. 2011). PhyloCSF-Ψ_Emp_ instead uses the empirical distributions of PhyloCSF scores on carefully selected coding and non-coding regions of different lengths, in order to reduce deviation between the tails of the actual and approximate distributions that limits the ability to distinguish protein-coding regions when extremely high specificity is needed.

We found 220 second ORFs for which the PhyloCSF-Ψ_Emp_ score is more than 17.0, a threshold chosen to account for the low prior probability that a second ORF is in fact a readthrough region. We excluded any transcripts for which the first stop codon is present only in close relatives of *A. gambiae*, as these could be recent nonsense substitutions that would leave a downstream protein-coding signature without true readthrough. We then manually examined the alignment for each of the remaining transcripts and excluded any for which it was likely that the protein-coding signature is due to an alternative splicing event, translation initiation at a downstream ATG, or translation on the opposite strand. Finally, we used SECISearch3 (Mariotti et al. 2013) to find likely selenoproteins and excluded one candidate, AGAP000358, a homolog of the known *Drosophila* selenoprotein SelG. This resulted in a list of 187 likely readthrough candidates which we designated as our “preliminary list”.

We next used information about the identity and conservation of the first stop codon to expand our preliminary list of candidates. Candidate readthrough stop codons found by evolutionary signatures in *Drosophila* have a striking tendency to use the same stop codon in all species, perhaps because the three stop codons encode different amino acids when read through or modulate the readthrough rate, and to preferentially use TGA and, to a lesser extent, TAG (Jungreis et al. 2011). This is also true of the subset for which readthrough was observed in ribosomal profiling experiments (Dunn et al. 2013).

We observed a similar pattern among the *Anopheles* readthrough candidates in our preliminary list (Figure 2C). We defined a new score for a second ORF, Adjusted-Ψ_Emp_, that combines PhyloCSF-Ψ_Emp_ with a likelihood ratio for the choice and degree of conservation of the first stop codon, estimated using our preliminary list (see Methods). Because of limited alignment quality of the two most distantly-related *Anopheles* species (*A. darlingi* and *A. albimanus*), we computed scores both with and without these two species and used the higher of the two scores. We added to our list of candidates 92 transcripts having second ORFs whose Adjusted-Ψ_Emp_ is more than 17.0 and for which we could find no other likely explanation for the protein-coding signature, as described above.

Because there are many signals that a second ORF is protein-coding that are not accounted for by Adjusted-Ψ_Emp_, such as frame conservation, length of the second ORF, cytosine immediately 3’ of the first stop codon, synonymous substitutions at the second stop, and low coding potential after the second stop, we manually examined the alignments of moderately scoring second ORFs and added 40 candidates to our list whose Adjusted-Ψ_Emp_ scores are somewhat less than 17.0 but that seemed likely to be readthrough based on these additional factors (Supplemental Figure S7).

Next, we examined annotated *Anopheles* transcripts orthologous to 282 previously reported *D. melanogaster* readthrough candidates (Jungreis et al. 2011), and added 21 to our list that would have passed our previous checks had we used a lower score threshold, on the assumption that orthologs of readthrough stop codons are more likely to be readthrough than other stop codons. We refer to these as candidates “found using orthology”.

Finally, for transcripts in which there is another stop codon immediately 3’ of the annotated stop codon, we applied a similar procedure to the ORF immediately 3’ of that second stop codon, and found 13 candidates for readthrough of two adjacent stop codons, which we refer to as “double-stop readthrough” (Figure 1D). This is a special case of reading through two stop codons that are not necessarily adjacent codons, which we refer to as “double readthrough”. We are not aware of any previous predicted or experimentally observed cases of double-stop readthrough.

The result was our final list of 353 *A. gambiae* transcripts for which the most plausible explanation of the observed evolutionary signature is functional and evolutionarily-conserved stop codon readthrough, not associated with selenocysteine insertion, henceforth referred to as the “readthrough candidates” (Supplemental_Data_S1.txt).

We applied a similar procedure to the regions between the second and third stop codons of the readthrough candidates to find 35 candidates for double readthrough, including the 13 cases of double-stop readthrough. Finally, among our double readthrough candidates there are five that show clear evolutionary signatures of triple readthrough (Figure 1C), including two in which a single readthrough stop codon is followed by a double stop codon in some species (Supplemental Figure S10). Triple readthrough has been previously predicted for two *D. melanogaster* genes (Crosby et al. 2015).

Because Adjusted-Ψ_Emp_, is a log likelihood ratio, we can use it to estimate the false discovery rate given a prior probability that a transcript is readthrough. Of the 74% of readthrough candidates for which the second ORF has Adjusted-Ψ_Emp_ > 17, we estimate the false discovery rate is 11%, 8%, or 6% for a prior probability of 0.02, 0.03, or 0.04, respectively. Below, we will present evidence that more than 4% of annotated stop codons are readthrough, so the false discovery rate is lower than 6%.

While it is also possible that some of the second ORFs in our candidate list are partly coding due to an alternate splice variant or a downstream start site, our manual inspection was intended to exclude such cases so it is unlikely that there are many remaining. We cannot estimate a false discovery rate for the remaining 26% of readthrough candidates because they were included based on unquantified additional evidence.

In order to facilitate cross-clade comparisons of orthologous readthrough stop codons, we applied a similar procedure to *D. melanogaster* orthologs of our *A. gambiae* readthrough candidates and identified 51 *D. melanogaster* readthrough candidates that we had not previously reported (Supplemental Figure S8), including one candidate for double-stop readthrough (Supplemental Figure S9D). Six of these 51 have been predicted to be readthrough transcripts previously (Crosby et al. 2015). Combining these 51 with 282 reported in our 2011 paper gave us 333 *D. melanogaster* readthrough candidates to be used in our downstream analyses (Figure 2D, SupplementalDataS1.txt).

## Insights into readthrough evolution from mosquito-fly comparisons

In order to characterize the evolutionary dynamics of readthrough, we quantified the typical features of readthrough transcripts, compared candidates in *A. gambiae* to those in *D. melanogaster*, and compared candidates that have orthologs in the other species to those that do not. By comparing orthologs between the two clades, we can see evolutionary effects over a considerably longer time scale than are revealed by orthology within either clade (Figure 3A).

**Figure 3.**
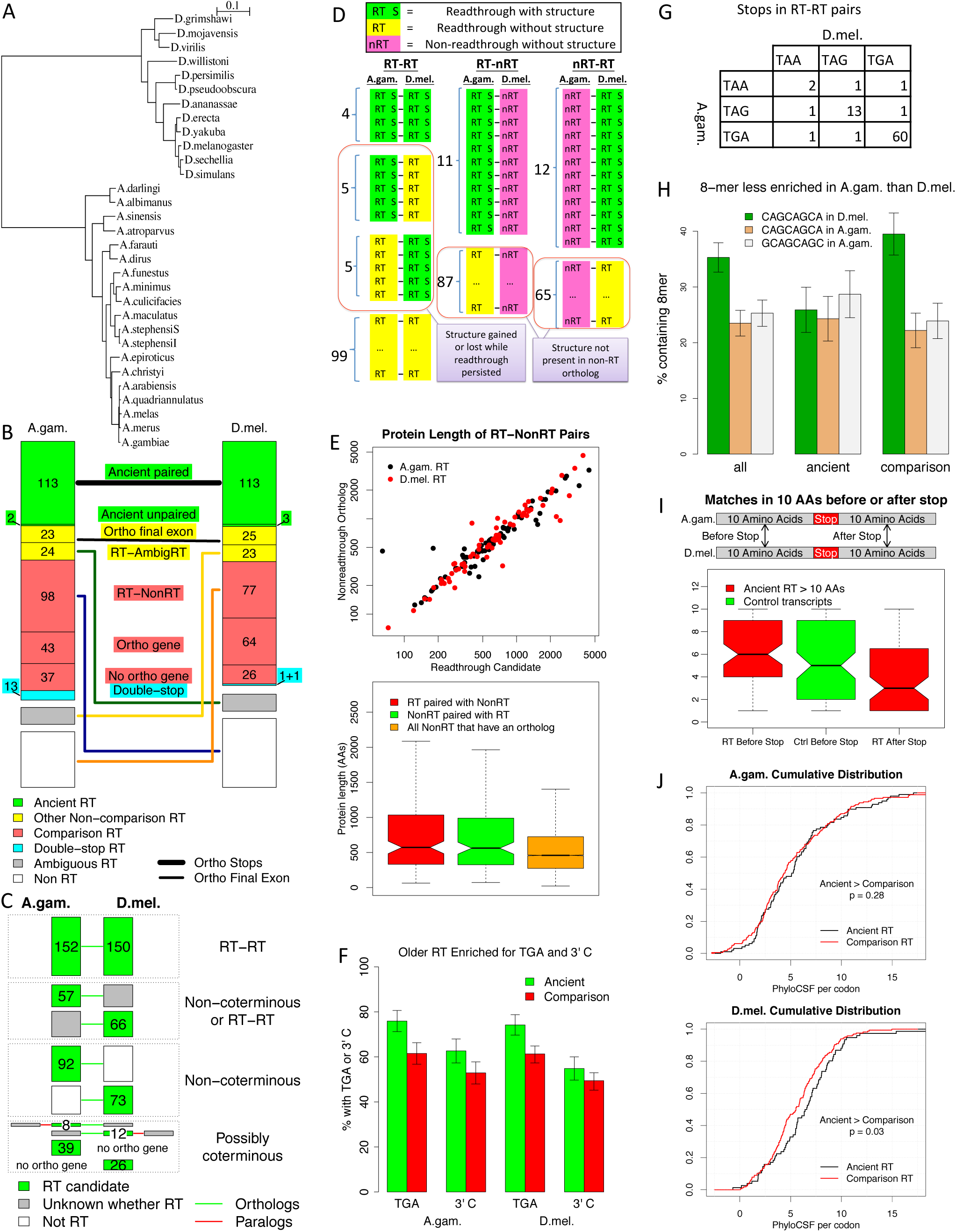
Mosquito-fly comparison provides insights into readthrough evolutionary dynamics. (A) Phylogenetic tree of 12 *Drosophila* and 19 *Anopheles* species estimated using concatenated protein sequence alignments of single-copy orthologs (the Mali-NIH and Pimperena strains of *A. gambiae* are not shown). The scale shows a genomic distance of 0.1 neutral substitutions per four-fold degenerate site. (B) Categorization and pairing based on orthology. Boxes quantify stop codons in *A. gambiae* (left) and *D. melanogaster* (right) that were used in our cross-clade comparisons. Filled boxes quantify readthrough candidates, color-coded by the category of orthology to the other species. Candidates likely to have been readthrough in the common ancestor, “ancient”, are green; candidates least likely to have been readthrough in the ancestor, “comparison”, are red; 13 double-stop readthrough in *A. gambiae*, one double-stop readthrough in *D. melanogaster*, and one readthrough in *D. melanogaster* found using orthology to a double-stop readthrough, all of which have been excluded from most of our comparisons, are cyan; and remaining readthrough candidates are yellow. Other boxes quantify stop codons that are clearly not conserved readthrough, even within their own clade, (white), or for which we could not make a confident determination of readthrough (gray), that are orthologous to readthrough candidates in the other species. The thick line joins boxes representing candidates in pairs having orthologous stop codons and the thin lines join boxes representing transcripts in pairs having orthologous final exons but not necessarily orthologous stop codons. Two readthrough candidates in *A. gambiae* and three in *D. melanogaster* are stop-orthologous to readthrough candidates in the other species but were excluded from some comparisons in order to make a one-to-one pairing. (C) Most differences in readthrough repertoires are not due to gene birth and death. Boxes classify and quantify the common and distinct portions of the readthrough gene repertoires of *A. gambiae* (left) and *D. melanogaster* (right), to determine which differences are associated with gene birth and death, “coterminous” events, and which are not. Homology relationships between readthrough candidate genes (green boxes) and non-readthrough genes (white) or genes of uncertain readthrough status (gray) are represented by green lines (orthologs) and red lines (paralogs). Some differences in readthrough gene repertoires might be due to coterminous events (bottom group of boxes), whereas others cannot be (second group of boxes from the bottom). There are other cases where we do not know if the repertoires are different but we know that if they are it is not due to coterminous events (third group of boxes from the bottom). At most 34% of the differences are due to coterminous events. (D) RNA structures in orthologous pairs. Three columns correspond to readthrough candidates in *A. gambiae* orthologous to readthrough candidates in *D. melanogaster*, readthrough in *A. gambiae* orthologous to non-readthrough in *D. melanogaster*, and non-readthrough in *A. gambiae* orthologous to readthrough in *D. melanogaster*. Readthrough candidates having a predicted structure are in green, readthrough candidates without a structure are in yellow, and non-readthrough orthologs, none of which have predicted structures, are in magenta. Among readthrough-readthrough pairs, nine have structures in *A. gambiae* and nine do in *D. melanogaster*, but only four have structures in both, implying that some structures are ancient whereas others have been gained or lost while readthrough persisted. None of the non-readthrough transcripts orthologous to readthrough candidates have structures, suggesting that the structures were not present for very long before readthrough appeared. (E) First ORF lengths for readthrough-non-readthrough pairs. Upper figure is a scatter plot showing the length of the first ORF of each *A. gambiae* readthrough candidate orthologous to a non-readthrough *D. melanogaster* transcript versus the first ORF length of its non-readthrough ortholog (black dots) and corresponding pairs of lengths for *D. melanogaster* readthrough candidates orthologous to non-readthrough *A. gambiae* transcripts (red dots). Lower figure is a box plot showing the first ORF lengths of those readthrough candidates in either species that are orthologous to a non-readthrough transcript in the other, the corresponding lengths of the paired non-readthrough transcripts, and the lengths of all non-readthrough transcripts in genes that have orthologs in the other species. There is almost no difference between the first ORF lengths of the readthrough candidates and their non-readthrough orthologs, but they are generally larger than the other non-readthrough transcripts, implying that longer genes are more likely to become readthrough rather than that genes tend to get longer after becoming readthrough. (F) Older readthrough are more likely to use TGA and C. The first stop codon is TGA in a significantly larger fraction of our ancient readthrough candidates (green) than of the readthrough candidates in our comparison group (red), in both *A. gambiae* (first column) and *D. melanogaster* (third column). The base immediately 3’ of the stop codon is C in a higher fraction of ancient readthrough candidates than of readthrough candidates in our comparison group (second and fourth columns), though the difference has limited statistical significance. Both comparisons were performed using a subset of readthrough candidates that were classified as readthrough without depending on the choice of stop codon. Error bars show standard error of mean. (G) Stop codon usage in ancient readthrough pairs. Table indicates the number of ancient readthrough candidates having specified stop codons in *A. gambiae* and *D. melanogaster*. The dearth of pairs having a TGA stop codon in one species and not the other (only 4 pairs) implies that the increased prevalence of TGA stop codons among ancient readthrough candidates is likely due to loss of readthrough among TAA and TAG stop codons, rather than conversion of TAA or TAG stop codons to TGA. (H) Enriched 8-mer. Chart shows the fraction of readthrough candidates that include the 8-mer CAGCAGCA in the second ORF or the final 250 nucleotides of the first ORF, in *D. melanogaster* (green) and *A. gambiae* (tan), as well as the corresponding fractions for the ancient and comparison subsets. Error bars show standard error of mean. This is the most prevalent 8-mer in these regions, and in our previous work we found it to be highly enriched among readthrough candidates even after correcting for GC content and other confounders. Also shown is the fraction among the same regions of *A. gambiae* candidates containing the 8-mer GCAGCAGC (white), which is the 8-mer that occurs in the most *A. gambiae* candidates although it is not the most prevalent when considering multiple occurrences in the same candidate. The CAGCAGCA 8-mer is highly enriched among readthrough candidates in each species, but is significantly more enriched in *D. melanogaster* than in *A. gambiae*. The difference between the two species is concentrated among the readthrough candidates in the comparison group, with almost no difference between the two species when considering only ancient readthrough, suggesting that the difference is due to an increased prevalence of the 8-mer in genes that have become readthrough in *Drosophila* since the lineages diverged, rather than due to genes that were readthrough in the ancestor gaining the 8-mer in the *Drosophila* lineage. (I) Readthrough regions under less purifying selection than other coding regions. Box plot showing the number of matching amino acids when aligning the 10 amino acids before or after the first stop codon with the corresponding region of the orthologous transcript in the other clade for readthrough-readthrough orthologous pairs whose readthrough regions are at least 10 amino acids long in each clade (red) and before the stop codon for orthologous pairs of control transcripts (green). The number of matches is significantly less for the readthrough regions than for the the regions before the first stop, whether comparing to the corresponding readthrough transcripts or to the control transcripts, implying that readthrough regions have been under less purifying selection at the amino acid level than other coding regions. (J) Ancient readthrough have higher PhyloCSF. Cumulative distributions of PhyloCSF per codon for the readthrough regions of ancient readthrough candidates (black) and candidates in the comparison group (red), for *A. gambiae* (upper panel) and *D. melanogaster* (lower panel). Scores are higher for ancient candidates, suggesting that older readthrough regions are under greater purifying selection at the amino acid level.

First, we verified that our *Anopheles* readthrough candidates have similar group properties to those previously reported for *Drosophila* (Jungreis et al. 2011). Specifically, there is a strong tendency for the first stop codon to be TGA, and for the base 3’ of the stop codon to be cytosine (C), both of which are known to increase translational leakage (Supplemental Figure S1); the first stop codon is highly conserved (Figure 2C); predicted conserved RNA structures are highly enriched in the 100 nucleotides 3’ of the first stop codon (9% of readthrough candidates versus fewer than 1% of non-readthrough transcripts); readthrough candidate genes tend to have longer first-ORF coding sequence, and tend to have more and longer introns (Supplemental Figure S2A-C); and the 8-mer CAGCAGCA is highly enriched within the second ORF and the 250 nucleotides 5’ of the first stop codon (23% of readthrough candidates versus 8% of non-readthrough transcripts).

Next, we determined pairs of readthrough candidates in the two species whose stop codons are orthologous, using pairs of Diptera-level *A. gambiae - D. melanogaster* orthologs from OrthoDB version 7 (Waterhouse et al. 2013). Many of these genes have alternative splice variants containing different stop codons, so for each pair of orthologous genes we identified which pairs of transcripts, if any, have orthology in the final exon or portion of an exon 5’ of the annotated stop codon, which we will refer to as having “orthologous final exons”. Among those, we looked for pairs for which we could detect orthology immediately 5’ of the first stop codon, which we will refer to as having “orthologous stop codons” or “stop-orthologous”, to exclude cases where the stop codon had moved in one clade due to a nonsense substitution or frameshift.

We found that 115 of our *A. gambiae* readthrough candidates are stop-orthologous to one or more *D. melanogaster* candidates, and 116 *D. melanogaster* candidates are stop-orthologous to one or more *A. gambiae* candidates (in a few cases several paralogous candidates are orthologous to the same candidate in the other species, Supplemental Figure S9A). For some of these pairs of orthologous readthrough stop codons, readthrough could have evolved independently along the two lineages from a non-readthrough ancestral stop codon, however we estimate that the number of cases of such convergent evolution is only around 7 (Supplemental Text S1). Consequently, we would expect that for almost all pairs of stop-orthologous readthrough candidates the ancestral stop codon in the common ancestor of *Drosophila* and *Anopheles* was readthrough, hence we refer to them as “ancient”.

Some special cases of orthologous readthrough candidates are shown in Supplemental Figure S9, namely four-way homology of two pairs of alternative splice variants (B), a double-stop readthrough candidate orthologous to a single-stop candidate (C), and a double-stop readthrough candidate orthologous to another double-stop candidate (D).

To understand how readthrough evolved within the two clades for genes that were already readthrough in the common ancestor, we selected a unique representative in each species for each set of many-to-one orthologs to obtain a set of 113 pairs of stop-orthologous readthrough candidates that we could use for cross-species comparisons (Supplemental Figure S9A,B). In each species we also defined a “comparison” group of readthrough candidates least likely to have been readthrough in the common ancestor, by excluding from the complete list of candidates any whose final exon is orthologous to a readthrough candidate, even if we had not classified the stop codons as orthologous, or to a transcript that we had not classified as readthrough but that we could not be certain was not readthrough, which we refer to as, “ambiguous readthrough”. That left 178 and 167 candidates in the *A. gambiae* and *D. melanogaster* comparison groups, respectively. We have no way to know whether these comparison candidates were readthrough in the common ancestor, because readthrough could have been gained in one clade or lost in the other, but our expectation is that the set is highly enriched for candidates that did not exist in the ancestor, so differences between candidates that were readthrough in the ancestor and ones that were not are likely to be detected by comparing our ancient and comparison groups. The classification of orthologs can be found in Supplemental_Data_S1.txt. We also defined a set of readthrough-non-readthrough pairs by taking the 98 *A. gambiae* and 77 *D. melanogaster* readthrough candidates in the comparison group whose final exons are orthologous to final exons of transcripts in the other clade that are unambiguously not conserved readthrough because of frame shifts in the second ORF or poor conservation of the second stop codon. The orthology classification is summarized in Figure 3B.

For each of the group properties of readthrough candidates previously identified, we report our findings from various comparisons of *A. gambiae* to *D. melanogaster*, ancient group to comparison group, and readthrough candidates to their non-readthrough orthologs in readthrough-non-readthrough pairs. In some cases we compared restricted subsets of candidates to avoid biases introduced by the curation process. In particular, we excluded candidates found using orthology from comparisons related to PhyloCSF because we used a lower score threshold for such candidates. Also, for comparisons related to stop codon choice and conservation, we defined a subset, “unbiased by stop codon”, that avoids biases introduced by the way some candidates were identified. In most cases we report results with double-stop readthrough candidates excluded because we only searched for these in *A. gambiae* and because of other possible biases, but we verified that including them would not affect any of the conclusions. Also, in most cases we report results for pairs having orthologous stop codons, but we verified that the conclusions remained valid if we included all pairs having orthologous final exons, even those we had not classified as having orthologous stop codons.

## Most readthrough birth and death is not due to gene birth and death

As a first application of our orthology classification, we investigated the dynamics of readthrough birth and death. Does readthrough tend to arise soon after a gene is born and then last for the full lifespan of the gene? Or can readthrough appear long after the gene matures or disappear while the gene persists? We will refer to the birth of readthrough soon after the birth of its gene or loss of readthrough only upon the death of its gene as “coterminous” readthrough events, whereas readthrough birth in an old gene or readthrough death before the death of its gene are “non-coterminous”. If readthrough birth and death are largely coterminous, we would expect differences in the readthrough repertoires of *A. gambiae* and *D. melanogaster* to be primarily due to genes that have arisen in one species or been lost in the other since the speciation event, whereas otherwise we would expect many ancestral genes surviving in both lineages to exhibit readthrough in one species and not the other.

To resolve the question, we obtained bounds on the number of each type (Figure 3C). Let N be the number of differences in the readthrough gene repertoires of the two species that resulted from non-coterminous events and C be the number due to coterminous events or to a combination of the two. We looked at the level of gene rather than transcript, because orthology between genes can be determined more reliably. We only considered genes of readthrough candidates, including double-stop readthrough candidates, and their homologs, since we have not identified other readthrough genes.

There are three ways that a coterminous readthrough event could lead to a difference in the readthrough repertoires of the two species: a readthrough gene arose de novo in one lineage; a gene that was readthrough in the ancestor was lost in one lineage; or a non-readthrough gene in the ancestor duplicated in one lineage and the new gene quickly became readthrough. (We have ignored more complicated scenarios, such as gene duplication followed by readthrough genesis in the duplicate and loss of the parent gene, as we would expect such combinations of rare events to be exceedingly rare.) In the first two cases there would be a readthrough gene in one species having no orthologous gene in the other species, whereas in the third case there would be a readthrough gene in one species having a non-readthrough paralog in the same species and a non-readthrough ortholog in the other species. The number of readthrough candidate genes satisfying one of these two conditions is thus an upper bound for the number of coterminous readthrough events among our readthrough candidates. We would not expect this bound to be sharp, since those same conditions can also have arisen through non-coterminous events. There are 65 readthrough candidate genes (39 in *A gambiae* and 26 in *D. melanogaster)* that have no orthologous gene in the other species, and there are 20 readthrough candidate genes (8 in *A. gambiae* and 12 in *D. melanogaster)* for which we found a Diptera-level paralog in the same species that is not a readthrough candidate gene and for which we did not find a readthrough ortholog in the other species. Thus at most 85 of the differences we found in the readthrough repertoires of the two species are due to coterminous events, 0 ≤ C ≤ 85.

On the other hand, there are 165 readthrough candidate genes (92 in *A gambiae* and 73 in *D. melanogaster)* that have no Diptera-level paralog (or all of whose paralogs are readthrough candidates) and whose final exon is orthologous to the final exon of a transcript in the other species that we classified as definitely not conserved readthrough (and is not also orthologous to the final exon of a transcript we classified as readthrough or ambiguous readthrough). For each of these, the difference between the two species must have arisen through a non-coterminous event. There are also 123 readthrough candidate genes (57 in *A. gambiae* and 66 in *D. melanogaster)* that have no non-readthrough paralog and for which either we found an ortholog that we classified as ambiguous readthrough or we could not identify a transcript with orthologous final exon; each of these might or might not be orthologous to a (non-candidate) readthrough gene but we can be sure that if it is not then the difference is due to a non-coterminous event. Since any of the 85 differences that could be coterminous might instead have been non-coterminous, we have 165 ≤ N ≤ 165 + 123 + 85 = 373

Thus, the fraction of differences in the readthrough gene repertoires that are due to coterminous events, C / (N + C) is at most 85 / (85 + 165) = 34%. The actual fraction is probably much lower because we do not expect our upper and lower bounds to be sharp.

Our conclusion is that most of the time readthrough arises long after the birth of the gene, or is lost before the death of the gene. We are unable to distinguish between these two possibilities, but finding the readthrough gene catalog of an outgroup species might enable such a determination in the future.

## RNA structures can be gained or lost while readthrough persists

We next used RNAz to predict conserved RNA structures in the 100 nt regions 3’ of readthrough stop codons. We had previously found a strong enrichment for such structures in windows of that size 3’ of *D. melanogaster* candidate readthrough stop codons (Jungreis et al. 2011), and such a structure has been found to trigger readthrough in the *Drosophila hdc* gene (Steneberg and Samakovlis 2001). RNAz combines predictions of thermodynamic stability and evolutionary conservation to make more robust predictions of RNA structures than either alone (Gruber et al. 2010).

We predicted RNA structures in 9% (33) our *A gambiae* readthrough candidates and 10% (34) of our *D. melanogaster* readthrough candidates compared to fewer than 1% of other transcripts (p < 1.0e-9).

We had previously found that the distribution of first stop codons among those readthrough candidates in *D. melanogaster* that have a predicted structure is significantly different from the distribution among candidates that do not (Fisher’s exact p = 0.0006) with more TAG and fewer TGA stop codons in the former, and speculated that a leaky stop codon context might not be necessary for readthrough in the presence of an RNA structure (Jungreis et al. 2011). However, among our *A. gambiae* readthrough candidates these distributions are not significantly different (p = 0.18, Supplemental Figure S4).

Among the 113 pairs of readthrough candidates having orthologous stop codons, nine have a predicted structure in *A. gambiae* and nine in *D. melanogaster* (Figure 3D). Four have predicted structures in both species, whereas the expected number if the presence of structures in the two species were independent is less than 1.0, suggesting that some of the structures were present in the common ancestor (p = 0.006). There is clear homology between stem loops near the 5’ ends of the predicted structures in AGAP007646-RA and FBtr0110970 (Supplemental Figure S3). Other than that, we see no obvious similarity between the predicted structures in the two species in each of these four pairs, offering the possibility that it is the presence of a stable structure that is functional rather than particular features of that structure.

The 113 pairs of stop-orthologous readthrough candidates include five having a predicted structure only in *A. gambiae* and another five having a predicted structure only in *D. melanogaster*. To determine if these mismatches were due to threshold effects or to misclassification of non-readthrough transcripts as readthrough candidates, we applied RNAz to 63 windows of various lengths and offsets on either side of the stop codon, and also reexamined the evidence for readthrough in each of these ten pairs. In at least three of the ten pairs, the evolutionary evidence of readthrough is unambiguously positive in both transcripts, the evidence that the stop codons are orthologous is strong, RNAz found a strong signal for a conserved RNA structure in one member of the pair, and RNAz did not find any signal for a conserved RNA structure in any of the 63 windows in the other member of the pair (AGAP004119-RA, FBtr0300330; AGAP005737-RA, FBtr0076636; and AGAP006528-RA, FBtr0075318). These three pairs show that in some cases an RNA structure can appear and undergo purifying selection long after readthrough had been established, or that an RNA structure can be lost while readthrough is maintained. We cannot distinguish between these two possibilities since we do not know whether the structures were present in the ancestor.

To learn more about the relative evolutionary timing of readthrough and structure formation, we looked for structures in the non-readthrough transcripts of our readthrough-non-readthrough ortholog pairs. Among the 98 *A. gambiae* and 77 *D. melanogaster* readthrough candidates paired with non-readthrough orthologs, RNAz predicted a structure in the 100 bases 3’ of the stop codon of 11 and 12, respectively, of the readthrough candidates, whereas it did not predict a structure in that window for any of the 175 non-readthrough orthologs. Among the 23 non-readthrough transcripts paired with a readthrough candidate that has a structure, the were only three for which RNAz predicted a structure in even one of the other 62 windows near the stop codon, and those could be false positives in light of the large number of windows tested. We conclude that for all or almost all readthrough candidates having structures that have gained readthrough since the two clades split, the structure was not present in the ancestor, and for all or almost all readthrough candidates for which both readthrough and a structure were present in the ancestor but readthrough was lost in one of the two clades, the structure was also lost in that clade. This implies that the structures are generally formed either after or shortly before readthrough is gained, and are lost either before or soon after readthrough is lost, since otherwise we would expect to see structures in many of the non-readthrough orthologs.

## Readthrough genes were long before they were readthrough

In our earlier work on *Drosophila*, we had observed that readthrough candidate genes were much longer than non-readthrough genes by many measures, however we were unable to make any inferences about causality (Jungreis et al. 2011). In order to explore this question we investigated gene lengths in ortholog pairs that are readthrough in only one of the two clades. Because many *A. gambiae* UTRs are not annotated, we restricted our investigations to three measures of gene length that do not include the UTRs, namely, the length of the spliced coding region (first ORF), the number of exons in the coding region, and the mean length of an intron within the coding region of each transcript that has at least one such intron.

First, we verified that by all three measures readthrough candidates are much longer than non-readthrough transcripts (rank sum p < 0.002 in each case; Supplemental Figure S2A-C). Then, for each orthologous pair among our readthrough-non-readthrough pairs, we compared the length of the readthrough candidate transcript to that of its non-readthrough ortholog, combining the two clades for greater statistical power.

We found that first ORF lengths of readthrough candidates are almost identical to those of their non-readthrough orthologs (Pearson correlation = 0.94, Figure 3E), but much larger than those of non-readthrough transcripts that have non-readthrough orthologs in the other species (rank sum p =.0004). This rules out the hypothesis that the transition to readthrough is associated with a lengthening of the first ORF and instead favors the alternative hypothesis that genes that already have a long first ORF are more likely to become readthrough.

Comparisons of intron length and number of exons between readthrough candidates and their non-readthrough orthologs were not conclusive, perhaps confounded by differential intron loss and shortening of introns in the two clades (Supplemental Figure S2D,E).

## Readthrough is more likely to be lost at TAA and TAG stop codons

We next compared the prevalence of TGA first stop codon and 3’ base C among readthrough candidates in our ancient and comparison groups, restricting our attention to our subset of candidates unbiased by stop codon, in order to determine if there is an age-dependence for these prevalences (Figure 3F). Use of TGA as the first stop codon is significantly more prevalent among ancient readthrough candidates than among readthrough candidates in the comparison group (75.9% versus 62.1% in *A. gambiae*, 74.2% versus 61.1% in *D. melanogaster*, two-sided p = 0.057 and 0.041, respectively). Similarly, the occurrence of cytosine as the base immediately 3’ of the first stop codon is more prevalent among ancient readthrough candidates than among readthrough candidates in our comparison group (62.7% versus 52.4% in *A. gambiae*, 54.8% versus 49.1% in *D. melanogaster*), though with limited statistical significance (two-sided p = 0.182 in *A. gambiae* and 0.438 in *D. melanogaster*).

By comparing stop codons in the two clades, we find that the most plausible explanation for the enrichment of TGA stop codons among ancient readthrough transcripts is that readthrough was more likely to be lost if the readthrough stop codon was TAA or TAG than if it was TGA. In principle, there are two other possible explanations. First, it could be that a larger fraction of readthrough stop codons were TGA in the ancestor than in extant lineages. However, that seems unlikely because the fraction is almost the same in the two lineages, and it would have had to change to that same value independently in both. Second, it could be that many readthrough stop codons that were TAA or TAG in the ancestor changed to TGA in the current lineages. Since such a conversion would occur independently in the two lineages, if it were common we would expect to find many ortholog pairs in which one clade had TAA or TAG and the other had TGA, but this is not what we find: Among our 81 readthrough ortholog pairs unbiased by stop codon there are only 4 that are TAA or TAG in one clade and TGA in the other (Figure 3G), and in each of these at least one member of the pair has a short second ORF and could have been misclassified as readthrough.

### *Anopheles* readthrough are less enriched for CAGCAGCA than *Drosophila*

We next investigated the 8-mer CAGCAGCA, which we had previously found to be highly enriched among the *D. melanogaster* readthrough candidates in the regions extending from 250 nucleotides 5’ of the first stop codon until the second stop codon (Jungreis et al. 2011). We first verified that CAGCAGCA is the most common 8-mer in these regions, both in our expanded list of *D. melanogaster* readthrough candidates and in our *A. gambiae* readthrough candidates, occurring 500 times in the former and 369 times among the latter.

This 8-mer occurs in 35.3% of *D. melanogaster* regions but only 23.5% of *A. gambiae* regions, and the difference is significant, even after adjusting for the slightly longer regions in *D. melanogaster* (two-sided p = 0.0084; Figure 3H).

Although CAGCAGCA is the most frequent 8-mer in the *A. gambiae* regions, the related 8-mer GCAGCAGC occurs in more regions (25.3%). The fraction of *D. melanogaster* regions containing CAGCAGCA and the fraction of *A. gambiae* regions containing GCAGCAGC are also significantly different (two-sided p = 0.035).

The increased frequency of CAGCAGCA among *D. melanogaster* readthrough candidates compared to *A. gambiae* is concentrated in the clade-specific candidates. In fact, the fraction of *ancient* readthrough candidates containing this 8-mer is almost the same in the two species, 24.3% in *A. gambiae* and 25.9% in *D. melanogaster*, whereas the difference is exaggerated in the comparison group of clade-specific candidates, 22.5% in *A. gambiae* and 39.5% in *D. melanogaster*. Among the ancient readthrough candidates, there is a modest but significant correlation between the presence of the 8-mer in the two orthologs (r = 39.8%, p = 6.5E-6).

The concentration of the *D. melanogaster* excess among clade-specific candidates tells us something about the arrow of causality. This excess might be due to the 8-mer causing readthrough, or both being caused by some other condition, but it cannot be due to readthrough increasing the prevalence of the 8-mer, since the latter would have increased the presence of the 8-mer among the ancient *D. melanogaster* readthrough candidates as well as clade-specific ones.

## Readthrough regions diverge faster than first ORFs

We next investigated how quickly readthrough region sequences diverge compared to other coding regions.

First, to quantify within-clade purifying selection, for each of our readthrough candidates in *A. gambiae* and *D. melanogaster* we computed two measures of protein-coding potential, PhyloCSF and Z curve score, for the readthrough region, the same-sized coding region at the end of the first ORF, and the non-coding third ORF (excluding double readthrough candidates). The Z curve score provides a single-species measure of protein-coding potential using mono-, di-, and tri-nucleotide frequencies (Gao and Zhang 2004). Similar comparisons have been performed previously for our earlier set of *D. melanogaster* readthrough candidates (Jungreis et al. 2011; Dunn et al. 2013). We found that in both clades, both PhyloCSF and Z curve scores of readthrough regions were intermediate between those of coding first ORFs and non-coding third ORFs, indicating that readthrough regions have been under weaker within-clade purifying selection for protein-coding features than other protein-coding regions (Supplemental Figure S5A-D).

We next compared the two readthrough regions in each pair of ancient readthrough candidates to understand divergence across the two clades. For many of these pairs, the two readthrough regions have quite different length (Pearson correlation 0.74, Supplemental Figure S5E), suggesting that in some cases readthrough regions can remain functional despite large changes in length.

For many pairs of orthologous readthrough regions no relationship between the amino acid sequences was visually apparent, suggesting that the extensions were under less selective constraint than other coding regions since the time the two clades diverged. To quantify this, for each pair of stop-orthologous readthrough candidates having readthrough regions at least 10 codons long in each species, we aligned the first 10 amino acids of the readthrough regions in the two species and counted the number of matching amino acids. To see how this compared to amino acid conservation in other coding regions, we first compared these counts to the corresponding counts for the 10 amino acids just before the first stop codon of these pairs and found that the number of matches is significantly lower for the readthrough regions. However, that is not a fair comparison because the method we used to define orthologous stop codons introduced an upward bias to the amino acid conservation of the regions before the first stop codon. To address this, we also compared to a set of pairs of control transcripts that are likely to have orthologous stop codons but that are not biased towards higher amino acid conservation before the first stop codon (see Methods). We found that the number of matching amino acids in the first 10 amino acids of the readthrough regions of our orthologous readthrough pairs is significantly less than the number of matches in the final 10 amino acids of the first ORFs of our control transcripts (mean for readthrough regions = 3.9 matches, mean for control first ORF ends = 5.4 matches, one-sided rank-sum p = 0.002, Figure 3I). We have found that PhyloCSF scores tend to be lower near the ends of transcripts than in other parts of the transcript (unpublished), implying that they are under weaker purifying selection, so the difference between readthrough regions and typical coding regions is probably greater than is demonstrated by our comparison of readthrough regions to the final 10 amino acids.

We conclude that the amino acid sequences of the readthrough regions have been under weaker purifying selection than those of other coding regions.

## Ancient readthrough candidates have higher PhyloCSF scores

We next examined PhyloCSF scores as a proxy for determining whether within-clade purifying selection at the amino-acid level in readthrough regions has varied depending on how long a stop codon has been readthrough. For this comparison we excluded candidates that were found using orthology because that classification process introduced a bias towards lower PhyloCSF score.

We found that readthrough regions of ancient readthrough candidates have somewhat higher average PhyloCSF scores per codon than those of readthrough candidates in the comparison group (Figure 3J; mean 5.56 versus 5.30 in *Anopheles*, 6.48 versus 5.59 in *Drosophila*, rank sum p = 0.276 and p = 0.030, respectively; see Methods). This comparison is highly range-restricted because our list of candidates includes only those readthrough regions that have high PhyloCSF score, so the relatively low statistical significance in *Anopheles* could be due to limited statistical power.

A consequence of the bias towards higher PhyloCSF scores among ancient readthrough regions is that the readthrough transcripts that are not in our candidate list, and that therefore have lower PhyloCSF scores, are less likely to be ancient than our readthrough candidates are, even when we exclude candidates that were found using orthology (which are always ancient), and this is particularly true in *D. melanogaster* because of the more conservative threshold used in cataloging candidates in that species.

### There are over 600 readthrough stop codons in *A. gambiae* and 900 in *D. melanogaster*

We next estimated the number of readthrough stop codons in *A. gambiae* and in *D. melanogaster*, including ones that cannot be identified individually using PhyloCSF, by comparing the score distributions of second ORFs in three frames. In our earlier work, we had applied a similar technique to estimate that there were over 400 readthrough stop codons in *D. melanogaster* (Jungreis et al. 2011). Using improved techniques we can now bound the number more precisely and find that the actual number is much larger.

We define the second ORF in frames 1 and 2 to be the region starting 1 or 2 bases after the stop codon, respectively, and continuing until the next stop codon in that frame. We computed PhyloCSF-Ψ_Emp_ for the second ORFs in each of the three frames for every annotated stop codon, excluding ones for which the second ORF overlaps another annotated coding region or for which the alignment of the stop codon has inadequate branch length (Figure 4A). Readthrough would only cause a high score in frame 0, whereas other explanations such as an alternative splice variant with a 3’ splice site within the second ORF, translation start at a downstream ATG, overlap with an antisense coding region, and chance, could cause a high score in any of the three frames, and our earlier analysis in *D. melanogaster* found that the latter explanations do not show a bias towards frame 0 (Jungreis et al. 2011). Thus, any excess of high-scoring second ORFs in frame 0 is an indication of readthrough, and the area between the density curves provides an estimate for the number of readthrough stop codons.

**Figure 4.**
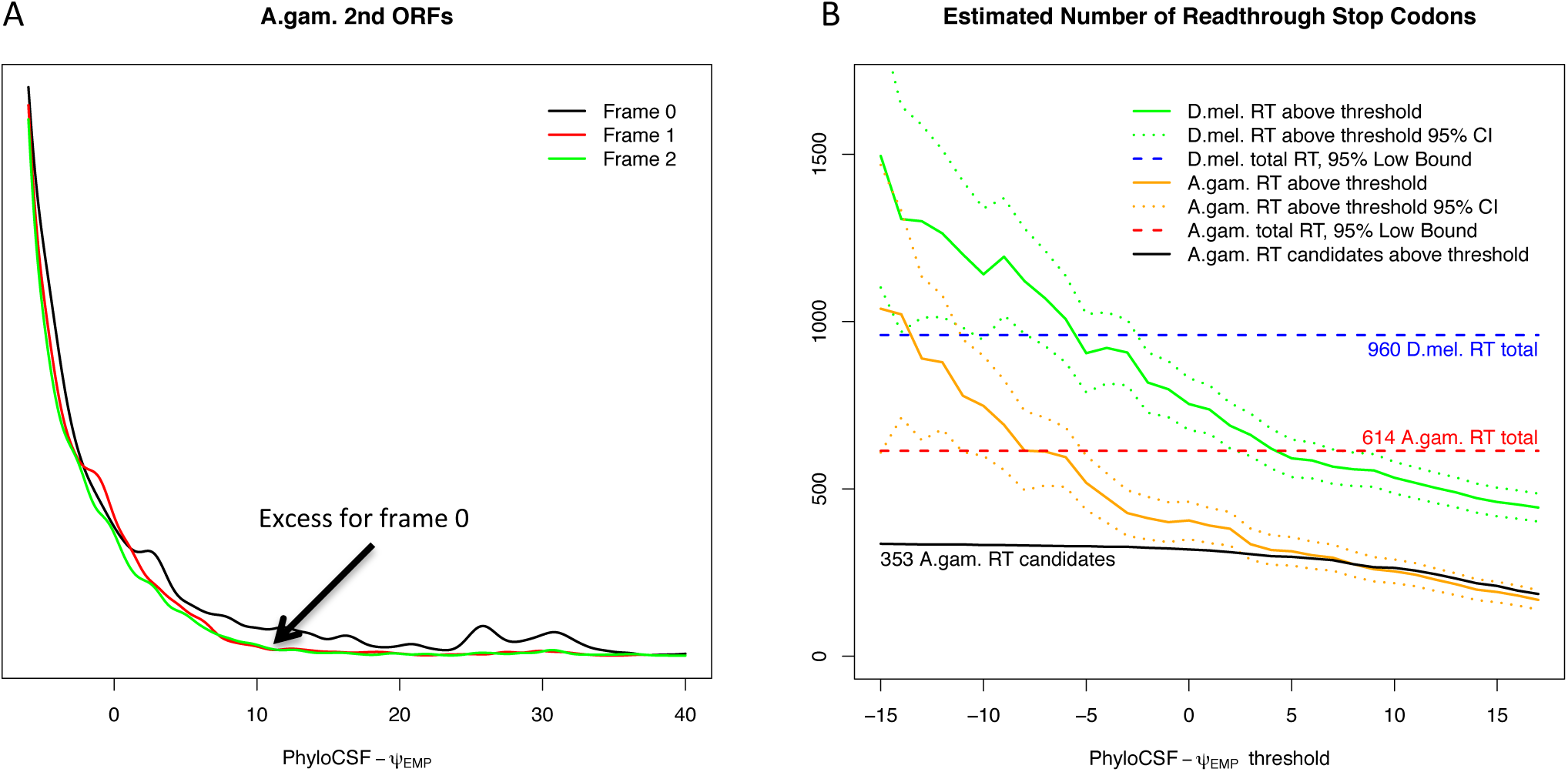
Estimating the number of readthrough stop codons. (A) Distribution of PhyloCSF-^v^P_E_mp scores of all regions starting 0, 1, and 2 bases after an annotated A *gambiae* stop codon (black, red, green, respectively) and continuing until the next stop codon in that frame, excluding ones that overlap an annotated coding region in any frame or whose alignment has inadequate branch length. Since readthrough second ORFs would have elevated score only in frame 0, whereas regions with high score due to other causes would be distributed among all three frames, the excess of high scoring regions in frame 0 allows us to estimate the number of readthrough stop codons, including ones that we cannot distinguish individually. (B) Graph showing, for each PhyloCSF-^v^P_E_mp score threshold, /, the estimated number of readthrough regions having score higher than *t,inA. gambiae* (orange) and *D. melanogaster* (green), with 95% confidence intervals (dotted curves), and the number of *A. gambiae* readthrough candidates whose readthrough regions have score higher than / (black curve). Also, 95% confidence lower bound for the total number of functional readthrough stop codons in *A. gambiae* (red dashed line) and *D. melanogaster* (blue dashed line). The estimated number of readthrough regions having score greater than 0 is 406 in *A. gambiae* and 754 in *D. melanogaster*, and the difference is unlikely to be due to differential annotation quality. The total numbers of functional readthrough regions of all scores are, with 95% confidence, at least 614 in *A. gambiae* and 960 inZ). *melanogaster*, which are much larger than the numbers of candidates reported individually. In *A. gambiae*, the number of readthrough candidates is close to the estimated number of readthrough stop codons for PhyloCSF-T’Emp ^>^ 50, indicating that our candidate list includes almost all high-scoring readthrough regions.

For every score threshold, we estimated the number of readthrough regions having PhyloCSF-Ψ_Emp_ score above the threshold, with 95% confidence intervals, by comparing the numbers of second ORFs in frames 0, 1, and 2 having score above the threshold (Figure 4B, and Methods). We estimate that there are 406 *A. gambiae* and 754 *D. melanogaster* readthrough regions having PhyloCSF-Ψ_Emp_ > 0 (95% CI 350-461 for *A. gambiae* and 676-831 for *D. melanogaster*). The estimated number of readthrough stop codons in *D. melanogaster* is much larger than the number in *A. gambiae*, and at least part of this difference is a true biological difference between the species because the difference is more than could be accounted for by the more comprehensive transcript annotations in *D. melanogaster* (see Methods).

The actual number of functional readthrough regions is larger than these estimates because some of them have PhyloCSF-Ψ_Emp_ ≤ 0. We estimated the number of these by looking at counts in the three frames having scores above a lower threshold, and using the distribution of coding scores to estimate the residual number of readthrough regions having score below that threshold. We used a score threshold of −10 which corresponds roughly to the median score of non-coding regions. We report a lower bound rather than an expected number because our estimate is highly sensitive to approximation error. We found that a 95% confidence lower bound for the number of readthrough stop codons is 614 in *A. gambiae* and 960 in *D. melanogaster* which is 5% or, respectively, 6% of all annotated stop codons. Thus, the total number of functional readthrough regions that have been under purifying selection at the amino acid level in a substantial portion of their respective genera is considerably larger than the 353 and 333, respectively, that we have catalogued here.

A substantial portion of these functional readthrough regions are short. When our calculations are restricted to second ORFs at least 10 codons long we find 95% confidence lower bounds of only 302 in *A. gambiae* and 460 in *D. melanogaster*, suggesting that more than half of functional readthrough regions are less than 10 codons long.

For score thresholds, *t* > 5.0, the number of our candidate *A. gambiae* readthrough regions having PhyloCSF-Ψ_Emp_ > *t* closely tracks our estimate for the total number of readthrough stop codons satisfying that condition (Figure 4B), suggesting that our list includes almost all readthrough regions having PhyloCSF-Ψ_Emp_ > 5.0. The remaining ones, having PhyloCSF-Ψ_Emp_ > 5.0, cannot be identified using this scoring method without increasing the false discovery rate.

## Two readthrough regions have peroxisomal targeting signals

In order to investigate possible functions of readthrough in our *Anopheles* readthrough candidates, we searched for peroxisomal targeting signals in the readthrough regions using the PTS1 Predictor server (Neuberger et al. 2003). While the function of most eukaryotic readthrough extensions is unknown, peroxisomal targeting signals have been predicted or experimentally observed in the readthrough extensions of several genes in human, fly, and yeast (Stiebler et al. 2014; Schueren et al. 2014; Dunn et al. 2013; Freitag et al. 2012).

We found a strong predicted peroxisomal targeting signal in the extension of AGAP010769 (PTS1 score 12.8, false positive probability 1.7e-4, Supplemental Figure S6A). The signal is present in all of its orthologs among the 21 *Anopheles* sequences, despite the presence of several radical amino acid substitutions among the final 12 amino acids, which is where the localization signal is thought to reside (Neuberger et al. 2003). AGAP010769 is the *A. gambiae* ortholog of *D. melanogaster* CG1969 (Supplemental Figure S6B), an N-acetyltransferase whose readthrough extension was previously predicted to contain a peroxisomal targeting signal (Dunn et al. 2013). The evolutionary conservation of the signal across the two clades despite the amino acid substitutions and two 3-base indels provides evidence that it is functional.

We also searched for peroxisomal targeting signals in the readthrough regions of the *D. melanogaster* readthrough candidates and found a predicted signal in transcript FBtr0082288 of Tetraspanin 86D (PTS1 score 8.9, false positive probability 6.3e-4, Supplemental Figure S6C). The signal is conserved as far as *D. kikkawai* but not in *D. ananassae* or beyond and the ortholog in *A. gambiae* does not appear to be readthrough. Tetraspanin 86D contains four transmembrane domains and is involved with nervous system development, border follicle cell migration, and positive regulation of Notch signaling pathway (Dornier et al. 2012; Hemler 2005).

### Readthrough is abundant in other *Anopheles* and *Drosophila* species but not in centipede

Recent publication of the genome sequence of the centipede Strigamia maritima (Chipman et al. 2014) permitted us to refine the phylogenetic extent of abundant readthrough.

In our previous paper, we described a method to estimate the number of functional readthrough stop codons in a species using only a single annotated genome (Jungreis et al. 2011). Much like the method we used above to estimate the number of readthrough transcripts in *A. gambiae* and *D. melanogaster*, the single-species method scores second ORFs in three frames, with a large excess in frame 0 indicating abundant readthrough; however, it assesses coding potential using the Z curve score, a lower-resolution discriminator than PhyloCSF but one that requires only a single annotated genome, and this only provided a conservative estimate of the number of functional readthrough regions at least 10 codons long and having positive Z curve score, which probably includes fewer than 25% of all functional readthrough regions (Supplemental Text S2). At the time, the test indicated the presence of dozens to hundreds of readthrough transcripts in all of the insects and the one crustacean tested, whereas all other species tested, including one arachnid, appeared to have considerably fewer, consistent with the fact that a search using PhyloCSF found only a handful of readthrough transcripts in human and *C. elegans*. At that time, we conjectured that the phenomenon of having hundreds of functional readthrough transcripts evolved along the Pancrustacea lineage after it split from the ancestors of arachnids.

We applied our 3-frame Z curve score test to 19 of the 21 *Anopheles* species (all except *A. gambiae* Mali-NIH and *A. gambiae* Pimperena, for which no annotations were available), all 12 *Drosophila* genomes, the *S. maritima* genome, and all of the genomes we had previously analyzed (Jungreis et al. 2011), using updated assemblies or annotations where available (versions, sources, and citations in Supplemental_Table_S1.pdf). For each genome, we computed both a maximum likelihood estimate and a 95% confidence lower bound for the number of functional readthrough regions at least 10 codons long and having positive Z curve score (Figure 5). It should be noted these can be underestimates in genomes with low sequencing quality or incomplete annotations (Supplemental Text S2). We defined “abundant readthrough” as having a maximum likelihood estimate more than 50, which, among the species we tested, is nearly equivalent to requiring that the 95% confidence lower bound is greater than 0.

**Figure 5.**
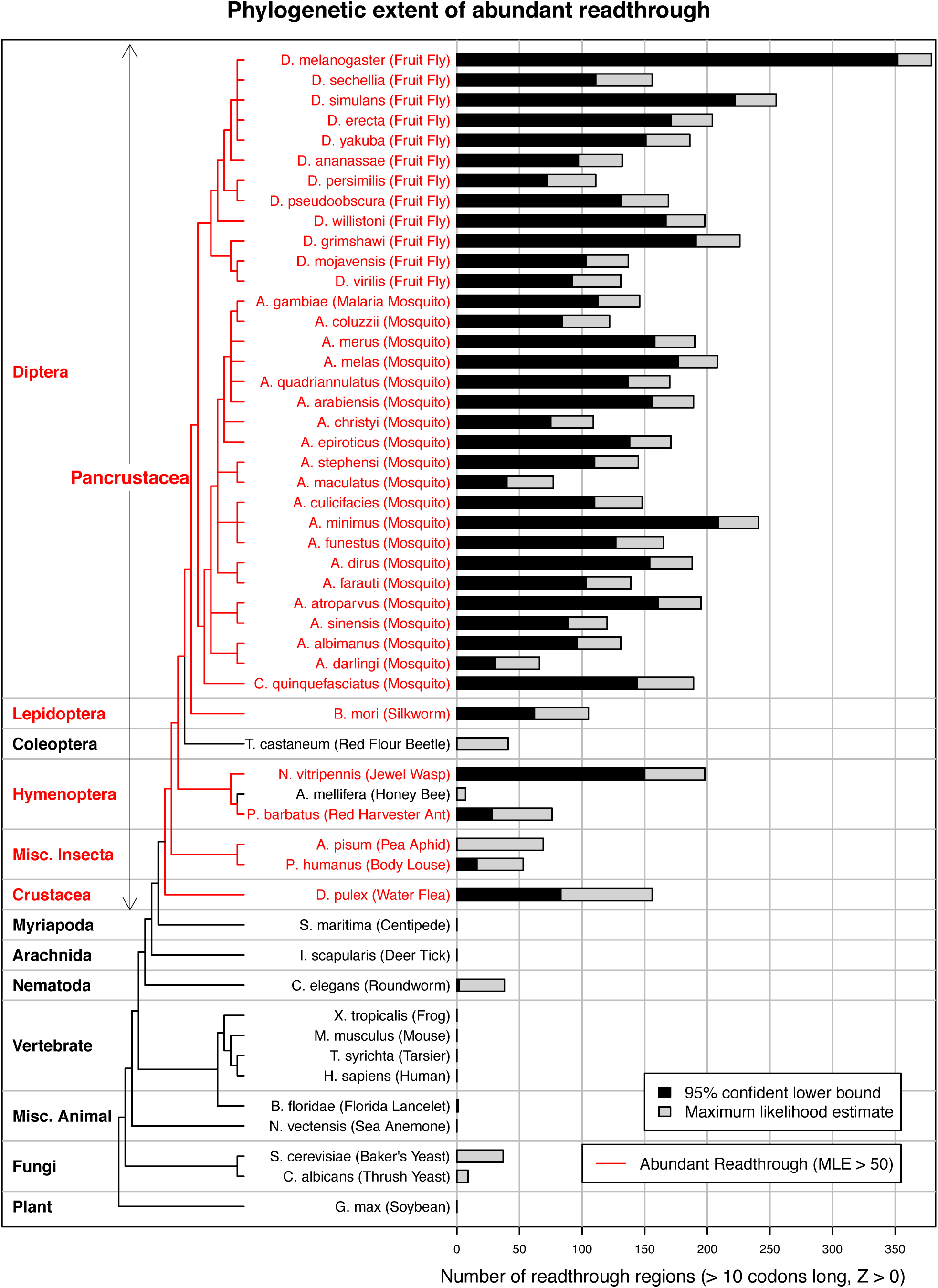
Estimated abundance of readthrough in 52 eukaryotic species. Estimate is calculated using single-species sequence-composition evidence quantified by Z curve scores for downstream ORFs in three frames to detect excess of positive scores in frame 0 associated with abundant readthrough. For each species, gray bar shows the maximum likelihood estimate of the number of functional readthrough transcripts among the subset of transcripts whose second ORFs are at least 10 codons long and have positive Z curve score, which probably includes fewer than one quarter of all functional readthrough transcripts, while black bar shows a 95% confidence lower bound. Tree shows phylogenetic relationships, with red branches indicating abundant readthrough, defined by maximum likelihood estimate greater than 50, which roughly corresponds to a 95% confidence lower bound greater than 0. Readthrough is abundant in all of the *Anopheles* and *Drosophila* species, most of the other insect species tested, and the crustacean, *D. pulex*, whereas none of the non-Pancrustacea species appear to have abundant readthrough, suggesting that it evolved in the Pancrustacea after they split from Myriapoda.

We found that all of the *Drosophila* and *Anopheles* genomes tested have abundant readthrough according to our definition, and in fact our 95% confidence lower bound exceeds 100 in almost all of those species. We suspect that the large excess in *D. melanogaster* as compared to the other *Drosophila* species is due to more complete annotations rather than to any biological difference. Among the other insects tested, *T. castaneum* and *A. mellifera* did not show abundant readthrough by our definition, though it is possible that our estimate is low due to incomplete annotations.

We found no frame-0 excess at all in the *S. maritima* genome, indicating few if any readthrough transcripts. This suggests that abundant readthrough evolved in the Pancrustacea after they split from Myriapoda (Figure 5), though we cannot rule out the possibility that abundant readthrough is present in other Myriapoda and was lost only in the *S. maritima* lineage, or, again, that our the test did not detect it due to incomplete annotations.

## Discussion

In this study, we found evolutionary signatures of functional, translational stop codon readthrough of 353 *A. gambiae* stop codons, supporting our earlier prediction that hundreds of genes in insect and crustacean species undergo functional stop codon readthrough.

We estimated that the number of stop codons undergoing functional readthrough is at least 600 (5%) in *A. gambiae* and 900 (6%) in *D. melanogaster*, enough to include one or more genes in most biological pathways. Since readthrough can have a major disease-relevant effect on the function of a protein, as illustrated by human VEGF-A in which readthrough converts an angiogenic protein to an antiangiogenic one, researchers will need to keep readthrough in mind when studying any aspect of insect or crustacean molecular biology. Our catalog of readthrough transcripts can be a starting point for efforts to characterize the function and regulation of the extended proteins.

Combining genomic data from multiple species in the *Anopheles* and *Drosophila* clades afforded several opportunities that were not available when data from only one clade was available. First, we used orthology to readthrough candidates in one clade in order to find readthrough candidates in the other clade that would have been missed otherwise, which resulted in 21 of our readthrough candidates in *A. gambiae* and an additional 45 *D. melanogaster* candidates that have not been previously reported. Second, we determined which properties are specific to the clade and which are more universal. We found two readthrough-related differences between the two clades, namely the larger estimated number of readthrough genes in *D. melanogaster* than *A. gambiae*, and the increased prevalence of the enriched CAGCAGCA motif in the *D. melanogaster* readthrough candidates than in the *A. gambiae* candidates. Finally, comparison of orthologs provided insights into the time scales and causal relationships that control the evolutionary dynamics of readthrough by giving us information about how long a gene has been readthrough and how long it has had some of the distinctive properties of readthrough genes. We found that readthrough does not usually appear soon after the birth of the gene and last for the life of the gene, but instead can appear or disappear during the life of the gene, suggesting that readthrough can be a mechanism for rapid adaptation to new environments; that associated RNA structures can be gained and lost while readthrough persists; that functional readthrough is more likely to be lost at TAA and TAG stop codons than at TGA stop codons; that longer non-readthrough proteins are more likely to become readthrough than shorter ones; and that older readthrough regions are under more selective constraint than newer ones, though both are under less constraint than other coding regions. Hypotheses about the function, mechanism, and regulation of readthrough can be tested against these observations.

The higher rate of amino acid evolution in readthrough regions than in other coding regions is consistent with the protein misfolding avoidance and protein misinteraction avoidance hypotheses, which posit that the protein sequence evolutionary rate is lower in proteins of higher abundance because of the greater deleterious effect of misfolding or misinteraction of such proteins (Zhang and Yang 2015). Since readthrough regions are translated at lower frequency than their first ORFs, the corresponding peptide extensions will have lower abundance and under these hypotheses would have higher evolutionary rate. On the other hand, it has been suggested that readthrough extensions might not provide any functional benefit, but rather that the slower-than-neutral evolutionary rates of their peptide sequences detected by PhyloCSF result simply from the need to avoid toxic misfolding or misinteraction when they are created due to occasional but unavoidable translational leakage at the stop codon (Zhang and Yang 2015). However, the high conservation of leaky stop codon contexts in most of the readthrough candidates militates against this explanation; indeed, if translation of the downstream region provides no benefit then stop codon contexts providing more robust termination would be preferred.

The prevalence of readthrough in insects offers a variety of models for investigating the mechanism and regulation of readthrough, which could lead to improved treatments for genetic diseases caused by nonsense mutations. Small molecules that induce readthrough have already been used to treat such diseases (Keeling et al. 2014; Dabrowski et al. 2015; Keeling and Bedwell 2010; Schmitz and Famulok 2007), and greater understanding of readthrough regulation could allow better targeting of such drugs to trigger readthrough of these nonsense mutations while fully allowing translation termination at other loci.

Efforts are underway to sequence and annotate the genomes of many insects and crustacea because of their important impact on disease and food production (i5K Consortium 2013). With readthrough so abundant in these species, it is important to recognize and annotate readthrough genes in order to complete the reference annotations of these genomes for used in studies to elucidate gene function. The representation of readthrough genes in *D. melanogaster* by FlyBase as alternative transcripts with longer CDS regions (Crosby et al. 2015) can serve as a model. Our techniques for finding readthrough genes should be applicable to any clade for which many genomes at the appropriate evolutionary distance have been sequenced and aligned, as is the case for bees (Kapheim et al. 2015) and ants (Simola et al. 2013). Our new techniques can also be used to more thoroughly search for *D. melanogaster* readthrough genes, as well as finding readthrough genes in the other species of the *Anopheles* and *Drosophila* clades.

## Methods

### Transcripts, Whole Genome Alignments, and Trees

We used version 4.2 of the *A. gambiae* genome assembly and annotations, obtained from VectorBase (Giraldo-Calderón et al. 2015). We used version 5.27 of the *D. melanogaster* genome assembly and annotations, obtained from flybase.org (Tweedie et al. 2009). Sources of the assemblies and annotations for the other species shown in Figure 5 are listed in Supplemental Table 1. We used the tree and divergences of 12 *Drosophila* species from (Stark et al. 2007) and the 12-flies subset of the 15-way dm3 insect alignments obtained from the UCSC Genome Browser (Kent et al. 2002; Kuhn et al. 2009).

The *Anopheles* whole genome multiple sequence alignments and phylogenetic tree were built using the 21 available *Anopheles* mosquito genome assemblies from VectorBase. The alignment building process is described in detail in (Neafsey et al. 2015). The set of assemblies includes *A. gambiae* PEST (Holt et al. 2002), *A. gambiae* Pimperena S form and *A. coluzzii* (formerly *A. gambiae* M form) (Lawniczak et al. 2010), the species sequenced as part of the *Anopheles* 16 Genomes Project (Neafsey et al. 2013), *A. darlingi* (Marinotti et al. 2013), and the Indian strain *A. stephensi* (Jiang et al. 2014). In summary: Multiple whole genome alignments of 21 available *Anopheles* assemblies were built using the MULTIZ feature of the Threaded-Blockset Aligner suite of tools (Blanchette et al. 2004), employing a similar approach to that used for other multi-species whole genome alignments such as those for 12 *Drosophila (Stark et al. 2007)* and 29 mammal (Lindblad-Toh et al. 2011) genomes. Before computing the alignments, repetitive regions within each of the input genome assemblies were masked. Assemblies were analysed using RepeatModeler (Smit and Hubley 2010) to produce repeat libraries that were then combined with known repeats from *A. gambiae* and retrieved from VectorBase, before being used to mask each genome assembly using RepeatMasker (Smit et al. 2014). The 21-species maximum likelihood phylogeny, required to guide the progressive alignment approach of MULTIZ, was estimated using RAxML (Stamatakis 2014) from the concatenated protein sequences of Genewise (Birney et al. 2004) gene predictions using Benchmarking Universal Single-Copy Orthologs (BUSCOs) from OrthoDB (Simão et al. 2015), and rooted with predictions from the genomes of *Aedes aegypti* (Nene et al. 2007) and *Culex quinquefaciatus* (Arensburger et al. 2010). The MULTIZ approach first runs all-against-all pairwise LASTZ alignments (default settings), followed by projections ensuring that the reference species is “single-coverage,” with projection steps guided by the species dendrogram to progressively combine the alignments.

The phylogenetic tree shown in Figure 3A was extracted from a 43-insects tree that included the 12 flies, 19 of the 21 *Anopheles* species (all except *A. gambiae* Mali-NIH and *A. gambiae* Pimperena), *G. morsitans*, *C. quinquefasciatus*, *A. aegypti*, *L. longipalpis*, *P. papatasi*, *D. plexippus*, *B. mori*, *T. castaneum*, *L. humile*, *A. mellifera*, *R. prolixus*, and *P. humanus*, and was built as follows. The maximum likelihood species phylogeny was estimated with RAxML (Stamatakis 2014)) using the PROTGAMMAJTT model on the concatenated protein sequence alignments of single-copy orthologs across all species, and rooted with the outgroups *P. humanus* and *R. prolixus*.

The non-metric phylogenetic tree shown in Figure 5 was extracted from Version 3 Draft synthetic tree of life, http://tree.opentreeoflife.org (Hinchliff et al. 2015).

### PhyloCSF and its derivates

PhyloCSF software, and parameters for the 12-flies alignment trained using *D. melanogaster* annotations, were obtained from github.com/mlin/PhyloCSF.git. We estimated empirical codon rate matrices for the 21-*Anopheles* alignments using the published algorithm (Lin et al. 2011) and the coding annotations of *A. gambiae*. However, we did not use these matrices and instead used the 12-flies rate matrices in *A. gambiae* as well as in *D. melanogaster* because this allowed better prediction of annotated coding regions in *A. gambiae* than the matrices estimated from *A. gambiae* annotations and alignments themselves, presumably because the *D. melanogaster* annotations are more accurate.

In what follows, we refer to the ratio of the branch length of the subtree of species present in the local alignment of a region to the branch length of the entire phylogenetic tree of the whole genome alignments as the “relative branch length” of the region.

For each of *A. gambiae* and *D. melanogaster* we defined PhyloCSF-Ψ_Emp_ of a region of length ***n*** codons having PhyloCSF score *Λ* as an approximation to the log of the ratio of the likelihood that a coding region of length ***n*** in that species would have a PhyloCSF score of *Λ* to the corresponding likelihood for a non-coding region, in units of decibans:

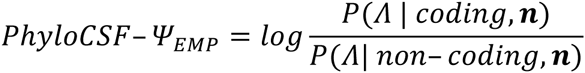

Rather than approximating the densities in this ratio with families of normal distributions, as was done to define PhyloCSF-Ψ (Lin et al. 2011), we used empirical distributions of scores of a training set of annotated coding and likely non-coding regions of various lengths in the corresponding species.

To maximize the specificity of PhyloCSF-Ψ_Emp_, it was critical to minimize the possibility that our non-coding training set included any regions that overlap possibly-unannotated coding regions. We also wanted to choose regions that would be as similar as possible to second ORFs, since those are the regions we intended to classify using PhyloCSF-Ψ_Emp_. To achieve these goals, we used regions at the 3’ ends of first ORFs of annotated coding regions and at the 5’ ends of corresponding *third* ORFs as our coding and non-coding training sets, respectively. We used third ORFs rather than second in order to avoid possible readthrough regions. We first compiled a list of all annotated coding transcripts whose final codon is a stop codon and for which neither the second ORF nor the 60 codons 3’ of the second stop codon overlap any annotated coding region in any frame on either strand or include any degenerate nucleotides (i.e., nucleotides reported as “N” in the genome assembly). For transcripts lacking an annotated 3’ UTR, or for which the 3’ UTR does not extend at least 60 codons beyond the second stop codon, we extended the transcript along the DNA strand without splicing. We compiled a list of coding and non-coding training regions by taking the first ***n*** codons 5’ of the first stop and the first ***n*** codons 3’ of the second stop, respectively, of each of these transcripts, for each value of ***n*** from 1 to 60, with the following exclusions: For both sets we excluded regions for which relative branch length is less than 0.1, since the PhyloCSF score on such regions is unreliable. From the coding set we excluded regions longer than the annotated first ORF. From the non-coding set we excluded regions longer than the third ORF. Also, to minimize the chance that a transcript undergoing double readthrough would be included in the non-coding training set, we excluded any regions for which the second ORF has PhyloCSF score ≥ 0 or the last 10 codons of the second ORF have PhyloCSF score ≥ 0, or for which the second ORF is too short or has too low relative branch length to rely on its PhyloCSF score (less than 10 codons or relative branch length less than 0.1). For *A. gambiae*, this left approximately 10,000 coding training regions of each length, whereas the number of non-coding training regions decreased from 5520 of length 1 to 385 of length 60. The corresponding numbers for *D. melanogaster* were 13,000, 6540, and 229, respectively.

For ***n*** ≤ **10** codons, we estimated the distribution of PhyloCSF scores of coding and non-coding regions of length ***n*** by applying the R language *density* function with default parameters to the scores of our training examples. For ***n*** > **10** codons, we did not have enough non-coding training examples to confidently estimate the density function in this way, so we instead scaled the density for regions of length 10 to match a mean and standard deviation specific to length ***n***. For **10** < ***n*** ≤ the mean and standard deviation were estimated from the scores of the training regions of that length; for ***n*** > **60** the mean and standard deviation were estimated by linear regression through the means and the logs of the standard deviations for 30 ≤ ***n*** ≤ 60. We limited PhyloCSF-Ψ_Emp_ scores to the range from −50 to 50, corresponding to likelihood ratios from 10^−5^ to 10^5^, because there were not enough training regions to define the tails of the empirical distributions beyond that point.

We defined Adjusted-Ψ_Emp_ of a second ORF as follows. For each aligned species, we determined if it has a stop codon aligned to the first stop codon of the transcript in the whole-genome alignments. We divided second ORFs into four categories, based on whether all such species have a TGA stop codon, all have a TAG, all have a TAA, or they do not all have the same stop codon. We defined Adjusted-Ψ_Emp_ of a second ORF with length ***n*** codons having PhyloCSF score *Λ* as:

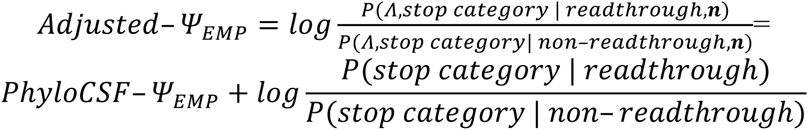

where the latter follows from the assumption that the probabilities of the stop categories are independent of the length and PhyloCSF score of the second ORF when conditioned upon the region being readthrough or non-readthrough, and considering readthrough second ORFs to be coding and non-readthrough as non-coding. We estimated the probabilities of the stop categories for readthrough and non-readthrough stop codons by counting them in our preliminary set of *A. gambiae* readthrough candidates and all other annotated stop codons having an aligned stop codon in at least 10 species, respectively. The resulting addends to PhyloCSF-Ψ_Emp_ were 8.5 for all TGA, 5.6 for all TAG, −2.8 for all TAA, and −15 for “mixed”. Because there were so few cases of the mixed category in our preliminary set, making the addend very sensitive to alignment errors, we flagged for manual examination any second ORFs having high PhyloCSF-Ψ_Emp_ but mixed stops, rather than accepting the number at face value. We used these same addends in both *A. gambiae* and *D. melanogaster*.

## Generating lists of candidates

For each annotated protein-coding nuclear transcript in *A. gambiae*, excluding transcripts whose final codon is not a stop codon, we computed the PhyloCSF-Ψ_Emp_ score for the second ORF, excluding the final stop codon. Among transcripts with identical second ORFs, we considered only one. For transcripts with no annotated 3’ UTR or for which the second ORF extended beyond the end of the annotated 3’ UTR, we defined the second ORF as continuing along the DNA strand beyond the end of the annotated transcript without splicing. We excluded second ORFs with relative branch length less than 0.1 as having inadequate branch length to compute a reliable PhyloCSF score.

For transcripts in which the genome assembly includes degenerate nucleotides in the second ORF, we truncated the second ORF at the first degenerate nucleotide. There is one such transcript, AGAP003849-RA, that was included in our final list of candidates because the 26 codons before the degenerate nucleotide provided adequate evidence of readthrough. We excluded this candidate from any analyses that depended on the entire second ORF.

To generate our preliminary list of *A. gambiae* readthrough candidates, we took second ORFs having PhyloCSF-Ψ_Emp_ ≥ 17.0 decibans, corresponding to a prior probability of being readthrough of approximately 0.02, which is roughly the fraction of *D. melanogaster* transcripts previously reported as readthrough candidates (Jungreis et al. 2011), and excluded ones for which we could find a more likely explanation for the high score than readthrough, according to the following criteria: If the relative branch length of the alignment of the stop codon is less than 0.1, we excluded that stop codon as a likely recent nonsense substitution. We generally considered the high PhyloCSF-Ψ_Emp_ to be explained by a coding exon of an alternative splice variant rather than readthrough if the second ORF contains an exon break or a predicted 3’-splice site with maximum entropy score (Yeo and Burge 2004) at least 4, and if PhyloCSF-Ψ_Emp_ < 0 for the region between the first stop codon and the exon break or predicted splice site, and we generally classified that transcript as being dicistronic rather than readthrough if PhyloCSF-Ψ_Emp_ < 0 for the region between the first stop codon and a fully conserved in-frame ATG or annotated start codon within the second ORF, though in a few borderline cases we adjusted that based on visual inspection of the alignment. If the PhyloCSF-Ψ_Emp_ score of the region on the opposite strand in the frame that shares the third codon position is higher than that of the second ORF, we consider the high PhyloCSF-Ψ_Emp_ of the second ORF to be explained by a coding region on the opposite strand, rather than by readthrough. We searched for SECIS elements in the 3’-UTRs of each of the remaining high-scoring second ORFs, extended to be at least 1000 nucleotides long, using SECISearch3 (Mariotti et al. 2013) with the default parameters, and as a result excluded AGAP000358, an ortholog of the known *Drosophila* selenoprotein SelG, from our candidates list as a likely selenoprotein; SECISearch3 found potential SECIS structures in two others, AGAP002233-RA and AGAP008574-RA, however these have low covariation scores, do not possess conserved adenines in the apical loop, and are not orthologous to any known selenoproteins, so we do not think they are real selenoproteins and did not exclude them from our list of readthrough candidates. Finally, we visually checked the alignments of the remaining second ORFs and excluded 10 for various reasons such as possible alignment errors.

In expanding our list of candidates using features indicative of coding regions that are not accounted for by Adjusted-Ψ_Emp_, we manually examined the alignments of all second ORFs whose Adjusted-Ψ_Emp_ score is between 5.0 and 17.0 and made a subjective assessment based on the following: high Adjusted-Ψ_Emp_ score of the initial 10 codons of the second ORF, second ORF more than 100 codons long, frame preserving indels, cytosine immediately 3’ of the first stop codon, high nucleotide conservation, synonymous substitutions in the second stop codon, and a sharp increase in nonsynonymous substitutions and frame-shifting indels after the second stop codon. We also included 2 exceptionally long second ORFs whose Adjusted-Ψ_Emp_ scores are less than 5.0, AGAP002296-RA and AGAP003059-RA.

We compiled the list of 282 *D. melanogaster* version 5.57 transcripts that we had previously reported as readthrough candidates by taking the 283 version 5.13 readthrough candidates reported in our previous paper (Jungreis et al. 2011) and for each one attempting to find a version 5.57 transcript having the same second ORF, or, if there is none, a version 5.57 transcript having the same stop codon. The stop codon of one of the 283 previously reported readthrough candidates, FBtr0078679, is no longer an annotated stop codon in version 5.57, so we did not include it in our analysis. The correspondence between previously reported version 5.13 transcripts and the version 5.57 transcripts used in the current study is indicated in Supplemental_Data_S1.txt. Six of the 51 *D. melanogaster* candidates that we identified using homology have been previously reported (Crosby et al. 2015), namely FBtr0345369, FBtr0084814, FBtr0345432, FBtr0079306, FBtr0330312, and FBtr0330733.

Any transcript whose final exon is homologous at the Diptera level to the final exon of a transcript already included in the readthrough candidate list for either species was added to the list if it satisfies the criteria described above except requiring PhyloCSF-Ψ_Emp_ ≥ 0 instead of PhyloCSF-Ψ_Emp_ ≥ 17.0, corresponding to a prior probability of readthrough of 0.5 rather than 0.02. One of the readthrough candidates identified in this way, FBtr0086577, was identified as a paralog of a readthrough candidate in the same species, FBtr0086599, while each of the others was identified as an ortholog of a readthrough candidate in the other species.

Our list of 353 *A. gambiae* candidates includes 28 that we had not included in (Neafsey et al. 2015). These were the 13 double-stop readthrough candidates, 10 additional orthologs of *D. melanogaster* readthrough candidates found using more refined criteria, and 5 others that were added based on careful examination of their alignments.

## Estimating false discovery rate

We estimated the false discovery rate for readthrough among the readthrough candidates having Adjusted-Ψ_Emp_ > 17, for a given prior probability of readthrough, ***pr***, as:

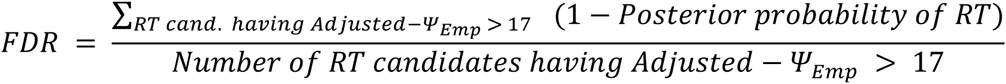

We used Bayes Theorem to get the posterior probability given the stop codon alignment, the second ORF length ***n***, and the PhyloCSF score, *Λ*, in terms of the prior, ***pr***, and the log likelihood ratio, which is approximated by Adjusted-Ψ_Emp_:

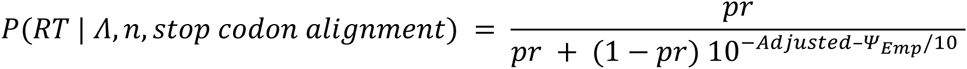

## Details of orthology classification

We considered two transcripts in homologous genes to have homologous final exons if there are four or more amino acid agreements in an alignment of the final ten amino acids of the final exon or if there are more amino acid agreements in an alignment of the final exon or portion of an exon 5’ of the stop codon of each than there are for 98.9% of pairs in the null model that we constructed, as described below. Alignments were calculated using the Needleman-Wunsch dynamic programming protein aligner NW-align (Yan et al. 2013). We made this relationship transitive by also considering two transcripts to have homologous final exons if both final exons are homologous to the final exon of some third transcript.

The cutoff of four amino acid agreements among the final ten amino acids was chosen because in a null model comparing alignments of the final ten amino acids of pairs of transcripts from non-homologous genes, 4.6% of pairs had three agreements and only 0.4% had four agreements. As a null model for the number of amino acid agreements in an alignment of the final exons of a pair of homologous transcripts, we counted amino acid agreements in an alignment of one or the other of those final exons with the final segment in a transcript of a non-homologous gene of the same length as the final exon of the other member of our pair. The cutoff of 98.9% corresponds to a false discovery rate of 0.02, and a local false discovery rate (Efron et al. 2001) of 0.3.

We considered two transcripts with homologous final exons to have homologous stop codons if their ultimate or penultimate amino acids agree, or if some third transcript is stop-codon-homologous to each.

The “unbiased by stop codon” subset of readthrough candidates was defined to be the 187 preliminary *A. gambiae* readthrough candidates, the 282 previously reported *D. melanogaster* readthrough candidates, and any non-double-stop readthrough candidates having PhyloCSF-Ψ_Emp_ > 0 whose final exon is homologous to the final exon of another readthrough candidate that is unbiased by stop codon. The 219 *A. gambiae* and 306 *D. melanogaster* readthrough candidates in our “unbiased by stop codon” subset are indicated in Supplemental_Data_S1.txt.

## Comparisons of readthrough properties

To calculate a p-value for the enrichment of the 8-mer CAGCAGCA in *D. melanogaster* readthrough candidates relative to *A. gambiae* readthrough candidates, we corrected for the different number and lengths of readthrough regions in the two species as follows: We chose 1000 random subsets of 331 of the 340 non-double-stop *A. gambiae* readthrough candidates. For each subset, we sorted the candidates by readthrough region length and paired them with the 331 non-double-stop *D. melanogaster* readthrough candidates, also sorted by readthrough region length. For each pair, we truncated the readthrough region of the longer one. We then counted the number of readthrough candidates in the resulting list for each species that contain the 8-mer in the 250 bases before the stop or in the (possibly truncated) readthrough region, and averaged these counts over the 1000 random subsets. Finally, we used the chi-squared test to calculate the two-sided p-value. The same method was used to calculate the p-value of the enrichment of CAGCAGCA in *D. melanogaster* compared to GCAGCAGC in *A. gambiae*. The one-sided p-value for the correlation between 8-mer presence in ancient readthrough pairs was calculated using the permutation test.

Structure prediction was performed using RNAz version 2.0pre, with the “-d” option, which compares to a null model that preserves dinucleotide frequencies, using the default cutoff of 0.5 for SVM RNA-class probability. Alignments were extracted for the the first stop codon and the next 97 nucleotides downstream of it using the 21-*Anopheles* alignments for *A. gambiae* and the 12-flies alignments for *D. melanogaster*, excluding species in which the alignment is not fully defined, and removing columns having gaps in all remaining species. To investigate the effect of thresholding on our classification, we also ran RNAz on alignments of windows of length 60, 80, 100, 125, 150, 175, and 200 nucleotides, starting at the first nucleotide of the first stop codon, or the nucleotides 20, 40, 60, or 80 nucleotides 3’ or 5’ of that nucleotide. For each of the 3 pairs we listed both of whose members are unambiguously readthrough and exactly one of which has a predicted structure, the second ORFs in each species have Adjusted-Ψ_Emp_ > 30, RNAz computed an SVM RNA-class probability of at least 0.98 in at least two windows for the one having a predicted structure, and computed an SVM RNA-class probability less than 0.06 in all windows for the other. The RNA structures in Supplemental Figure S3 were rendered using RNAplot from the Vienna RNA package version 2.1.8 (Lorenz et al. 2011).

The p-values in our comparisons of lengths of readthrough and non-readthrough candidates were computed using the one-sided Wilcoxon rank-sum test. For these comparisons, we did not exclude double-stop readthrough candidates because the analysis did not involve the second ORF. We excluded transcripts whose start codons are not ATG, since they are most likely truncated annotations of longer transcripts. We observed that genes that have Diptera-level OrthoDB orthologs in the other species tend to be longer than genes that do not, so when comparing coding sequence lengths of readthrough candidates having non-readthrough orthologs to lengths of non-readthrough transcripts, we restricted the latter to ones with Diptera-level OrthoDB orthologs in the other species, in order to eliminate the presence of an ortholog as a confounding factor in the comparison.

For comparing amino acid conservation in readthrough regions and other coding regions, we needed to find a set of pairs of control transcripts likely to have orthologous stop codons in a way that did not bias them towards higher amino acid conservation in the final 10 amino acids. We defined this control set to be all pairs of (not necessarily readthrough) orthologous genes in OrthoDB with no paralogs at the Diptera level, for which there is only one annotated transcript in each species, for which the coding sequence of that transcript lies within a single exon, and for which the the number of species in the multispecies alignment that have an aligned first stop codon is at least 11 in the *Anopheles* clade and at least 9 in the *Drosophila* clade (those being the minimum numbers of aligned stop codons among the readthrough pairs). The conclusion that the number of matches in the first 10 amino acids of the readthrough regions of our orthologous readthrough pairs is significantly less than number of matches in the final 10 amino acids of the first ORFs of our control transcripts remains true even if we do not require the coding regions to be single-exon (mean 4.5, p = 0.018), or do not require anything about the number of aligned stop codons (mean = 4.9, p = 0.013).

In comparing PhyloCSF scores of readthrough regions of ancient readthrough candidates to ones in the comparison group, we used PhyloCSF per codon rather than PhyloCSF-Ψ_Emp_ because the data points interpolated by the empirical distributions used to define the latter were too sparse to provide meaningful distinction among regions having PhyloCSF-Ψ_Emp_ much more than 17. We calculated the p-values for this comparison using a one-sided rather than a two-sided Wilcoxon rank-sum test because we had a prior expectation that the ancient ones would have higher PhyloCSF scores, due to the following logic: Our three-frame comparison showed that considerably fewer annotated *A. gambiae* stop codons are readthrough than *D. melanogaster* ones–only 69% as many even if we use the low end of the 95% confidence interval for *D. melanogaster* and even if we adjust the maximum likelihood estimate for *A. gambiae* to account for unannotated coding regions overlapping annotated stop codons (there is no need to adjust for unannotated stop codons because we are only considering annotated stop codons here). On the other hand the number of *D. melanogaster* readthrough stop codons that are orthologous to a readthrough stop codon in *A. gambiae* must be roughly equal to the number of *A. gambiae* stop codons that are orthologous to a readthrough stop codon in *D. melanogaster*. (They might not be exactly equal because several paralogous stop codons in one species can be orthologous to the same stop codon in the other; however, among our readthrough candidates there are very few for which this is true.) Consequently, the probability, p_D_, that a randomly chosen *D. melanogaster* readthrough stop codon is orthologous to a readthrough stop codon in *A gambiae* must be no more than around 69% of the corresponding probability in *A gambiae*, p_A_. However, our readthrough candidates are not randomly chosen readthrough stop codons, and could be more or less likely to have a readthrough ortholog than a randomly chosen readthrough stop codon. Let the probability that a *D. melanogaster* readthrough candidate has a readthrough ortholog be p_D_κ_D_ and the corresponding probability for *A gambiae* be p_A_κ_A_, where we exclude from consideration all candidates found by orthology since all of them have readthrough orthologs. Then we have p_D_ ≲ 0.69 p_A_, and by counting readthrough candidates that have orthologs we find that p_A_κ_A_ ~ 0.31 and p_D_κ_D_ ~ 0.27. Consequently, κ_A_ ≲ 0.79K_D_. Since the most noticeable difference between the *D. melanogaster* and *A gambiae* candidates is that the former have higher PhyloCSF-Emp scores due to more conservative curation criteria, it is natural to hypothesize that the reason κ_D_ is larger than κ_A_ is that readthrough regions having a higher PhyloCSF score are more likely to have a readthrough ortholog, or equivalently, that readthrough regions of ancient readthrough candidates tend to have higher PhyloCSF scores than other readthrough candidates.

## Estimating the number of readthrough regions using PhyloCSF-Ψ_Emp_

To estimate the number of readthrough stop codons in *D. melanogaster* and *A gambiae* using a 3-frames comparison of PhyloCSF-Ψ_Emp_ scores, we considered only nuclear transcripts for which the annotated coding sequence ends in a stop codon and the relative branch length of the alignment of the first stop codon is at least 0.1, choosing one representative transcript from each set of transcripts that share a stop codon. When counting second ORFs in each of the three frames having PhyloCSF-Ψ_Emp_ above some threshold, we considered only transcripts for which the second ORF in that frame is at least one codon long (not including the final stop codon) and does not include any degenerate nucleotide, the relative branch length of the alignment of the second ORF is at least 0.1, there is no annotated coding sequence that includes the first stop codon as a codon in that frame (for frames 1 and 2), there is no annotated coding sequence in any frame on the same strand that overlaps the second ORF but not the first stop codon, and there is no annotated coding sequence in any frame on the opposite strand that overlaps the second ORF (whether or not it overlaps the first stop codon). Let N0 be the number of second ORFs in frame 0 that satisfy these conditions.

To estimate the number of readthrough regions we modeled PhyloCSF-Ψ_Emp_ scores of readthrough regions as being drawn from a distribution *D*_1_, and scores of non-readthrough second ORFs drawn from a distribution *D*_2_ that is the same in all three frames. Let *f* be the fraction of frame-0 second ORFs that are readthrough. Then the scores of frame-0 second ORFs are modeled as being drawn from a mixture distribution *D*_3_ defined by:

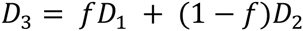

For any threshold, *t*, let *p*_*k*_(*t*) = *Prob*(*D*_*K*_ > *t*) and r(*t*) be the number of readthrough regions having score greater than *t*. Then,

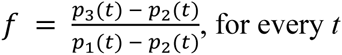

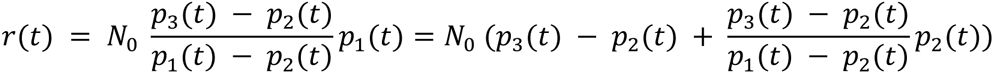

We have written the latter term to clarify the dependence on p_1_. For any *t*, we obtain a maximum likelihood estimate for p_3_(*t*) by counting second ORFs in frame 0 whose score is greater than *t*, and we estimate p_2_(*t*) similarly by counting second ORFs in frames 1 and 2; we obtain confidence bounds using a normal approximation to the binomial distribution. We do not know the score distribution for readthrough regions, so we estimate p_1_(t) using annotated coding regions; as we have seen earlier, coding regions tend to have somewhat higher PhyloCSF scores than readthrough regions, so this is likely to be an overestimate for p_1_, which gives us underestimates for *f* and *r*(*t*).

We estimate the total number of readthrough stop codons (not just those with score above some threshold) as:

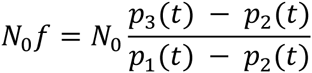

This would be the same for all values of *t* were it not for the approximations used to estimate p_1_, p_2_, and p_3_. Minimizing the approximation error requires choosing a value of *t* for which the probability that PhyloCSF-Ψ_Emp_ > *t* is similar for readthrough regions and other coding regions but substantially different for non-coding regions, the former because we are using the coding distribution as a surrogate for the readthrough region distribution and the latter to prevent catastrophic cancellation in the denominator. The former condition requires that *t* be less than the score of most readthrough regions, while the latter requires that *t* be more than the score of a substantial fraction of non-coding regions. As a suitable compromise between these opposing conditions, we chose *t* = −10, which is roughly the median score of non-coding regions.

To test if the more comprehensive annotations in *D. melanogaster* can fully explain the difference between *D. melanogaster* and *A. gambiae* of the estimated number of readthrough regions having PhyloCSF-Ψ_Emp_ > 0, we considered two kinds of missing annotations. First, the total number of annotated stop codons is 28% larger in *D. melanogaster*. If the two species in fact have the same number of stop codons and this difference is entirely due to some true *A. gambiae* stop codons not being annotated, then we would expect the real number of *A. gambiae* readthrough regions to be about 28% higher than our estimate. Second, when doing our three-frames comparison we excluded stop codons that are within an annotated coding region in another frame, because their second ORFs are more likely to have a positive score in frame 1 or 2 than in frame 0. There are many more such annotated overlaps in *D. melanogaster* than in *A. gambiae*, suggesting that *A. gambiae* probably has many *unannotated* overlaps. These probably inflate the counts of positive-scoring second ORFs in frames 1 or 2 and thus decrease our readthrough estimate below the actual number of readthrough stop codons. To correct for this, we estimated the number of such unannotated overlaps in *A. gambiae* as follows. For each of the sets of transcripts that were inputs to our calculation of N_0_, p_2_(t), and p_3_(t), we counted the number for which the 10 codons before the first stop have higher PhyloCSF-Ψ_Emp_ score in frame 1 or 2 than in frame 0, and for which the second ORF in that frame has PhyloCSF-Ψ_Emp_ > 0. We would expect many of the transcripts whose stop codons are within a coding region in another frame to satisfy this condition, but there would also be many false positives and false negatives. We estimated the false positive and false negative rates by determining what fraction of the stop codons satisfying the same conditions in *D. melanogaster* are within annotated coding regions in another frame. We then estimated the number of stop codons in *A. gambiae* overlapping (annotated or unannotated) coding regions in other frames by assuming that the false positive and false negative rates are the same in the two species. By subtracting these estimates of the *true* number of overlaps from the various counts in *A. gambiae*, rather than excluding only *annotated* overlaps, we corrected for the difference in this aspect of annotation quality between the two species. Doing this increased our estimate of the number of readthrough stop codons in *A. gambiae* having PhyloCSF-Ψ_Emp_ > 0 from 406 to 463. If we were to further increase this by 28% to account for the lower number of annotated stop codons in *A. gambiae* than in *D. melanogaster*, the resulting estimate of 592 readthrough stop codons in *A. gambiae* having PhyloCSF-Ψ_Emp_ > 0 is still well below the low end of the 95% confidence interval for the number of readthrough stop codons in *D. melanogaster* having PhyloCSF-Ψ_Emp_ > 0 (676). We obtained a similar result if, as an alternative to the above method, we estimated the number of readthrough stop codons in *A. gambiae* and *D. melanogaster* without excluding stop codons within annotated coding regions in other frames, in either species, which would inflate the estimate for the number of readthrough stop codons but do so similarly in both species. Thus, our conclusion that *D. melanogaster* has more readthrough genes than *A. gambiae* is unlikely to be an artifact of differences in annotation quality.

## Z curve score and single-species readthrough estimate

The Z curve 3-frames comparison test to estimate the number of readthrough stop codons in a genome was performed as described in (Jungreis et al. 2011). In brief, we trained the linear discriminant for the Z curve score in each species so that a score of 0 would be 50 times as likely for a coding region as for a non-coding region. We computed scores of second ORFs in three frames that were at least 10 codons long and did not overlap an annotated coding region in any frame and then calculated the number of second ORFs having positive score in frame 0 minus an average for the other two frames. We subtracted an estimate of the number of recent nonsense substitutions, which could lead to a false signal of readthrough, by scaling an estimate of the number of such mutations in *D. melanogaster* (17) by the total number of transcripts. We also subtracted an estimate of the number of sequencing errors that could similarly give a false signal of readthrough, obtained by finding orthologous regions in a related species that had a sense codon instead of a stop codon and adjusting the number of such regions by the fraction of simulated nonsense sequencing errors with downstream ORF having a positive Z curve score that could be detected by the same procedure. The related species used to find sequencing errors was *D. melanogaster* for all the insects, crustacea, and myriapoda, except the *Drosophilae*, for which we used *A. gambiae*; *T. urticae* for *I. scapularis*; *C. briggsae* for *C. elegans*; mouse for human and human for the other vertebrates; *S. cerevisiae* and *C. albicans* for each other; we did not subtract any estimate of sequencing errors for *G. max*, *B. floridae*, or *N. vectensis*, since the readthrough estimate was already 0. Reasons that the test is likely to underestimate the actual number are discussed in Supplemental Text S2).

## Data Access

Parameters for running PhyloCSF on *Anopheles* alignments (using the *Anopheles* tree and rate matrices trained on *Drosophila* alignments) have been added to github.com/mlin/PhyloCSF.git.

Supplemental_Data_S1.txt contains the list of readthrough candidates with coordinates, orthology information, links to CodAlignView, and other pertinent data.

Supplemental_Table_S1.pdf contains genome sources for the species in Figure 5.

Supplemental_Material.pdf contains Supplemental Text S1-S2 and Supplemental Figures S1-S10.

## Acknowledgements

We thank Marco Mariotti, Howie Waldman, Pouya Kheradpour, Eva M. Novoa, and Maxim Wolf for helpful discussions. IJ was supported by National Institutes of Health R01 HG004037 and GENCODE Wellcome Trust grant U41 HG007234. RMW was supported by Marie Curie International Outgoing Fellowship PIOF-GA-2011-303312.

### Author Contributions

CC, IJ, and MK designed the study, carried out the computational analysis and wrote the manuscript. RMW built the *Anopheles* alignments, delineated orthology and the species phylogeny, and helped write the manuscript. GF performed the search for peroxisomal targeting signals. ML supplied software for training PhyloCSF rate matrices.

